# Photoactivatable Large Stokes Shift Fluorophores for Multicolor Nanoscopy

**DOI:** 10.1101/2022.12.02.518850

**Authors:** Ilya Likhotkin, Richard Lincoln, Mariano L. Bossi, Alexey N. Butkevich, Stefan W. Hell

## Abstract

We designed caging-group-free photoactivatable live-cell permeant dyes with red fluorescence emission and ∼100 nm Stokes shifts based on a 1-vinyl-10-silaxanthone imine core structure. The proposed fluorophores undergo byproduct-free one- and two-photon activation, are suitable for multicolor fluorescence microscopy in fixed and living cells and are compatible with super-resolution techniques such as STED (stimulated emission depletion) and PALM (photoactivated localization microscopy). Use of photoactivatable labels for strain-promoted azide-alkyne cycloaddition and self-labeling protein tags (HaloTag, SNAP-tag), and duplexing of imaging channel with another large Stokes shift dye have been demonstrated.

---

Fluorescent dyes with large Stokes shifts are valuable tools in multicolor fluorescence microscopy, since they expand color multiplexing^1^ and allow shifting the detection window beyond the range of cell autofluorescence^2^ (<580 nm). These fluorophores typically have ≥80 nm difference between the excitation and emission band maxima for the lowest energy electronic transition (or ≥2000-4000 cm^-1^, depending on the spectral range). Red- and near-infrared (NIR) emitting fluorophores are particularly interesting,^3^ because they reduce phototoxicity and enable deeper optical penetration into the sample. Fluorophores with very large Stokes shift demonstrate zero overlap between absorption and emission spectra resulting in no self-quenching by reabsorption, and enable the quantitative interpretation of fluorescence signal irrespective of the label concentration.^4^

The growing usage of super-resolution light microscopy (nanoscopy) in routine biological imaging increases the demand for small-molecule photoactivatable or photoswitchable fluorescent probes.^5^ Unfortunately, there have been but singular reported examples of photoactivatable^6^ and photoconvertible^7^ large Stokes shift fluorophores, with none of them being suitable for tagging arbitrary proteins in living cells.

We have recently proposed a general synthetic strategy towards compact and biocompatible photoactivatable 1-vinyl-3,6-diaminoxanthone derivatives (PaX dyes^8^). Upon irradiation with UV or visible light, these compounds undergo rapid and clean intramolecular 6-*endo*-*trig* cyclization into the corresponding xanthylium fluorophores (Figure 1a) without generation of reactive electrophilic or radical intermediates and avoiding the need for photolabile protecting (caging) groups. Combining this idea with our earlier design of live cell-compatible large Stokes shift fluorescent probes,^9^ based on the pioneering works by Klán^10^ and Burgess,^11^ we have developed and report herein caging group-free photoactivatable fluorophores with Stokes shifts ≥100 nm.

**Figure 1.**
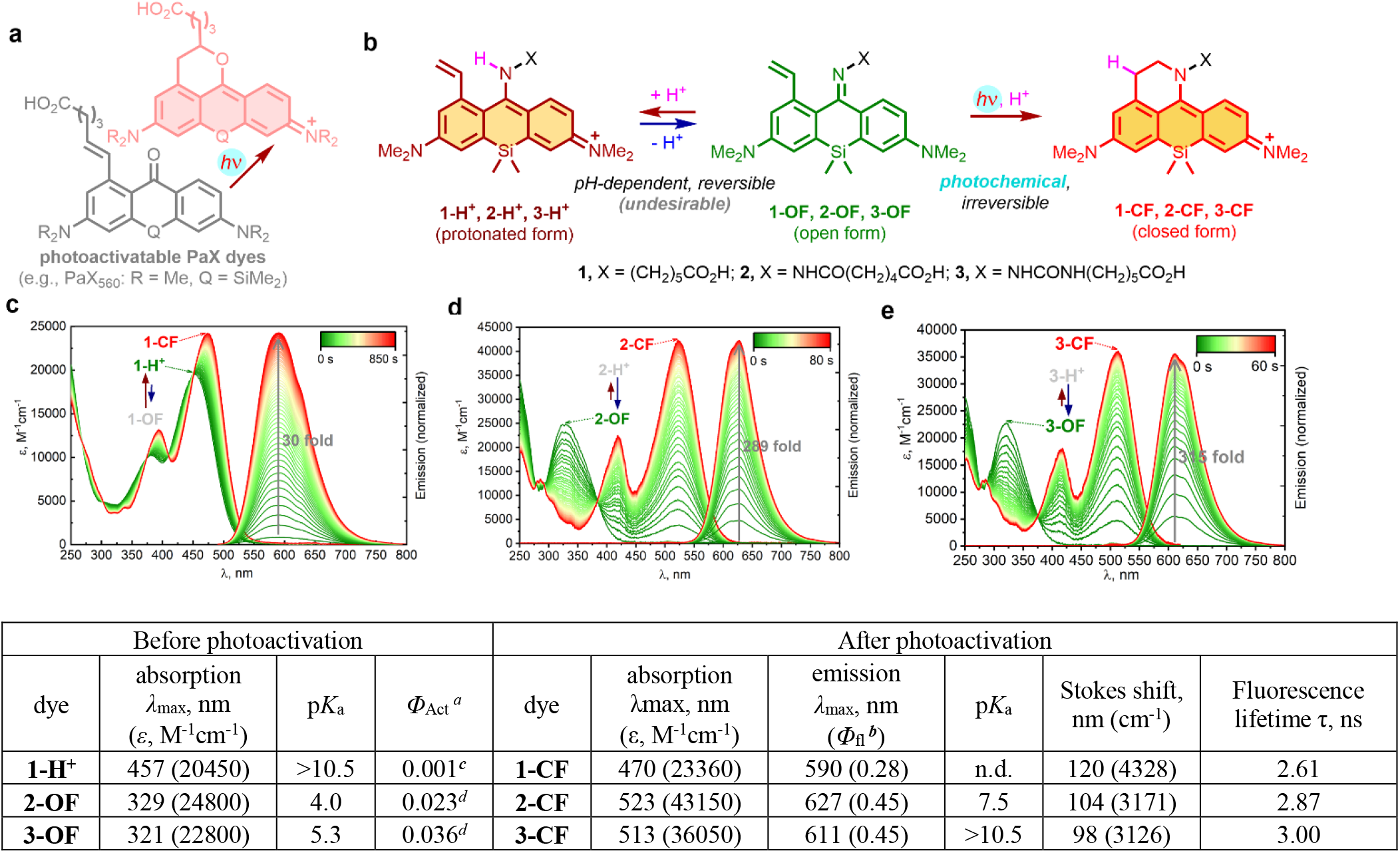
Photoactivatable large Stokes shift fluorescent dyes based on the 1-vinyl-10-silaxanthone imine core structure. (a) Photochemical generation of fluorophores from photoactivatable 1-vinylxanthone (PaX) dyes.^8^ (b) Reversible protonation of imines **1-OF**…**3-OF** vs. an irreversible photochemical ring closure. (c-e) Changes in absorption and fluorescence emission spectra of **1-OF** (c), **2-OF** (d) and **3-OF** (e) upon UV irradiation. ^*a*^ Photoactivation quantum yield; ^*b*^ fluorescence quantum yield; ^*c*^ irradiated at 405 nm; ^*d*^ irradiated at 365 nm.

In our initial attempt, an imino analogue **1** of the PaX_560_ core^8^ was prepared (Figure 1b). Even though this dye demonstrated sufficient fluorescence increase upon activation with 405 nm (or 455 nm) wavelength LED-light in aqueous media (30-fold in phosphate buffer at pH 7.0, Figure 1c) and the staining was specific, the fluorescence intensity before photoactivation in immunostained samples tested with several secondary antibodies was found unacceptably high (Figure S1), resulting in only two-fold emission enhancement upon UV irradiation. We hypothesized that switching from benzophenone imine to benzophenone oxime or hydrazone analogues should lower the contribution of the unwanted protonated state (e.g., **1-H**^**+**^) for the unactivated **OF** form of the dye. Following a brief *N*-substituent screening, an acyl hydrazone **2** and a semicarbazone **3** were identified as maintaining sufficiently large Stokes shift of the activated forms **2-CF, 3-CF** while showing clear preference for imine forms **2-OF, 3-OF** before photoactivation (Figure 1d,e) at physiologically relevant pH values. The dyes **2** and **3** demonstrated virtually indiscernible absorption and emission spectral profiles with one unit p*K*_a_ difference (Figure S2a,b), and underwent complete and byproduct-free photoactivation (Figure S3a,b). On studying the pH behavior of the model compounds, we noticed that the fluorescent closed form **2-CF** was losing its long-Stokes emission above pH > 7 (possibly due to deprotonation of the hydrazone NH; Figure S2c), while the photoactivated semicarbazone derivative **3-CF** remained pH-insensitive across the entire pH 3–10 range (Figure S2d).

We initiated our imaging studies by immunostaining the tubulin filaments in fixed COS-7 fibroblast-like cells with secondary antibodies labeled with reactive esters **2-NHS** or **3-NHS** (Figure 2a). Both labels could be activated in one-photon or two-photon mode with 405 nm or 800 nm activation laser, respectively (Figure S4, S5), and were found to be compatible with 775 nm pulsed STED light of relatively high peak intensity (∼60 MW/cm^2^), allowing to resolve the individual filaments with subdiffraction resolution (Figure 2b,c and Figure S6). However, the samples labeled with **2-NHS** demonstrated non-negligible initial fluorescence (24% for **2-NHS** vs. 5% for **3-NHS**) and required pre-bleaching of the label with 485 nm or 518 nm laser to approach zero background intensity levels before activation. For this reason, the dye **3** was preferred for single-molecule localization imaging. With the probe **3-Halo**, it was possible to resolve the circular shape of individual nuclear pores (outer diameter 107 nm^13^) in fixed U2OS cells stably expressing one of the nuclear pore complex proteins (Nup96) as a fusion with the HaloTag protein (a self-labeling modification of *Rhodococcus rhodochrous* chloroalkane dehalogenase DhaA^14^) using photoactivated localization microscopy (PALM; Figure S7).

**Figure 2.**
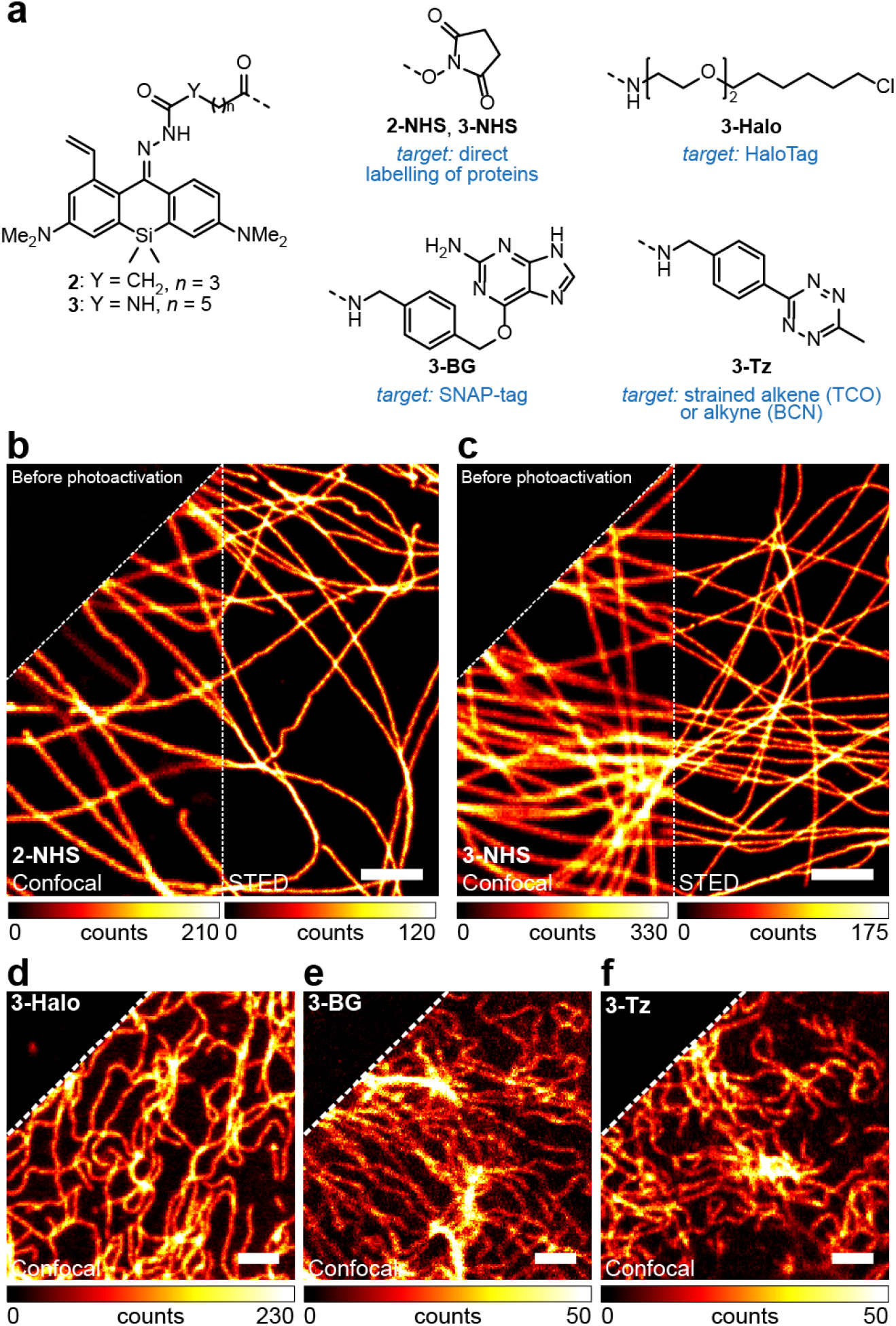
(a) Photoactivatable large Stokes shift fluorescent labels for fixed- (b,c) and live-cell (d-f) fluorescence imaging. (b,c) Confocal and STED images of tubulin filaments in fixed COS-7 cells labeled by indirect immunofluorescence with secondary antibodies tagged with **2-NHS** (b) or **3-NHS** (c). (d) Confocal image of U2OS cells stably expressing vimentin-HaloTag fusion protein labeled with **3-Halo** (200 nM). (e) Confocal image of U2OS cells stably expressing vimentin-SNAP-tag fusion protein labeled with **3-BG** (1 µM). (f) Confocal image of U2OS cells stably expressing vimentin-HaloTag fusion protein labeled with HTL-BCN (10 µM) followed by **3-Tz** (1 µM). In upper left corner of (b-f), the same images before photoactivation are shown. Scale bars: 2 µm.

To validate the selectivity of intracellular targeting with the ligands derived from photoactivatable large Stokes shift dye **3** in living cells, several labeling strategies have been explored. First, a vimentin-HaloTag fusion protein stably expressed in U2OS cells, genetically engineered using CRISPR-Cas9 technology (U2OS-Vim-Halo cells^15^), was labeled directly using **3-Halo** ligand. The indirect labeling was performed by pretreating the cells with excess bicyclo[6.1.0]non-4-yne (BCN) HaloTag ligand (HTL-BCN) followed by treatment with the tetrazine derivative **3-Tz**. Alternatively, living U2OS cells with stable expression of vimentin fused with SNAP-tag protein^15^ (an engineered human *O*^6^-alkylguanine-DNA alkyltransferase AGT^16^) were stained with *O*^6^-benzylguanine ligand **3-BG**. As illustrated by the Figure 2d-f, in all cases the bright and high-contrast selective staining of vimentin filaments has been achieved, with very little fluorescent background preceding the activation of the dye and no visible off-targeting in the activated samples. Beside these universally applicable self-labeling protein tags, selective targeting of **3** to lysosomes (Figure S8) was confirmed for a non-covalent Pepstatin A ligand (potent inhibitor of the ubiquitous lysosomal aspartic protease cathepsin D, binding to its active form^17^) by colocalization with the commercial *SiR-lysosome* probe.^18^

Relatively low sensitivity of the photoactivatable dyes **2**,**3** to visible light (>450 nm) permits channel duplexing with another large Stokes shift fluorophore sharing the same excitation wavelength and detection window. In our example, this scheme was realized in an immunostained sample by sequential imaging of the microtubules labeled with a moderately photostable coumarin dye DyLight 515-LS (excitation λ_max_ 515 nm, emission λ_max_ 650 nm) followed by photobleaching with the 485 nm excitation laser (Figure 3a,b). Photoactivation of **3**-labeled secondary antibodies used in indirect immunofluorescence labelling of TOMM20 protein was then achieved with 405 nm light and followed by confocal and STED imaging of mitochondria using identical instrument detection settings. This sequence provided a pseudo-two-color image for two large-Stokes fluorophores with overlapping absorption and emission spectra (Figure 3d-g).

**Figure 3.**
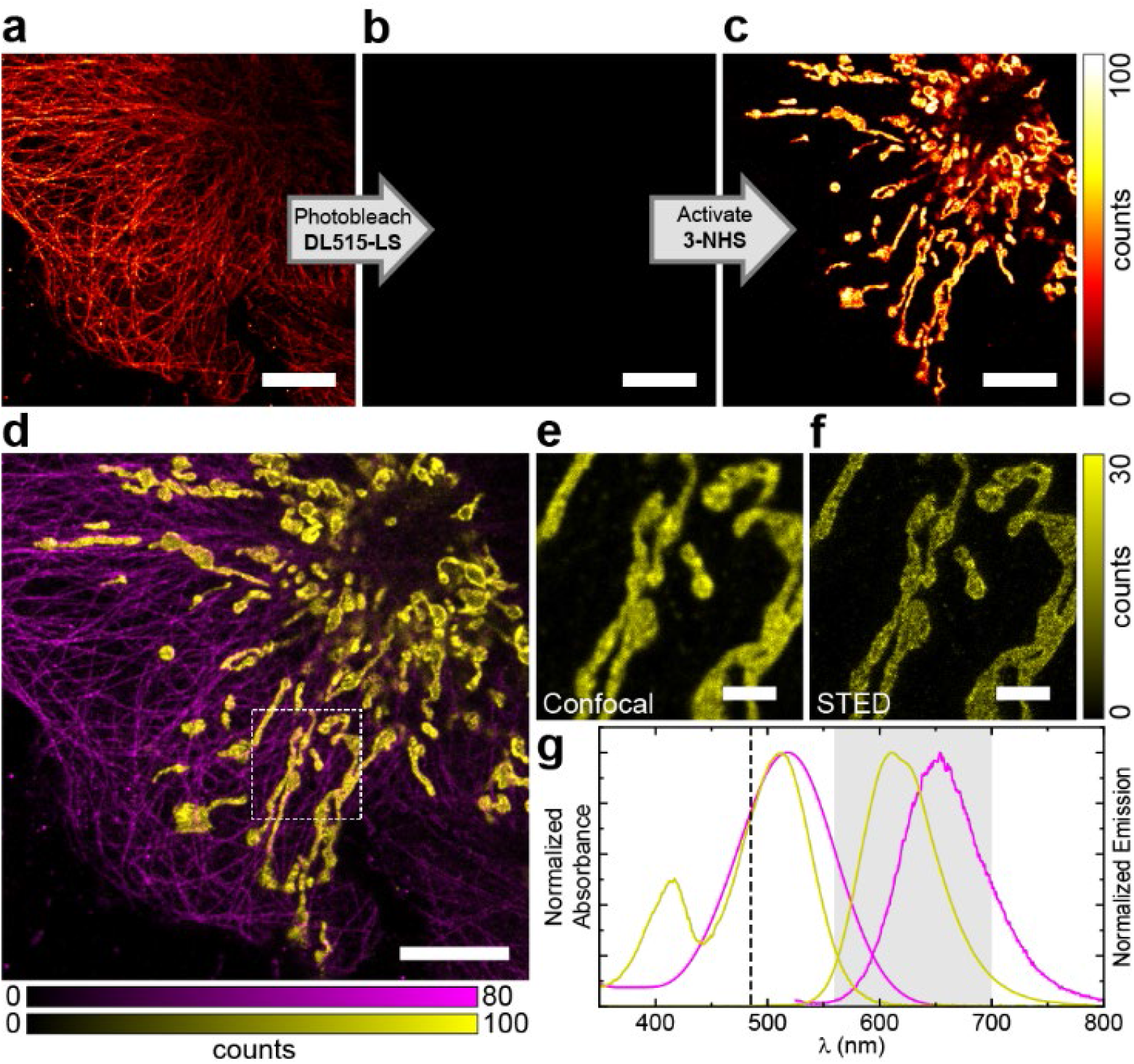
Channel duplexing with photoactivatable large Stokes shift dye **3**. (a-c) Confocal imaging of fixed COS-7 cells with microtubules labeled by indirect immunofluorescence with DyLight 515-LS NHS ester and mitochondria labeled with **3-NHS** before (a) and after (b) photobleaching of DyLight 515-LS and after (c) photoactivation of **3** by a 405 nm laser. (d) Combined pseudo-two-color image showing microtubules (magenta) and mitochondria (yellow) obtained by sequential imaging as shown in (a-c). (e,f) Confocal (e) and STED (f) images of mitochondria in the magnified region marked in (d). (g) Absorption and emission spectra of DyLight 515-LS (yellow) and **3** (magenta), the excitation laser line (485 nm, dashed line) and the detection window (560– 700 nm, grey); STED wavelength of 775 nm. Scale bars: 10 µm (a-d), 2 µm (e-f).

Finally, we demonstrated the use of photoactivatable large Stokes shift labels in four-color confocal imaging of fixed and living mammalian cells (Figure 4). In glutaraldehyde-fixed COS-7 cells, TOMM20 protein in mitochondria and vimentin in intermediate filaments were labeled by means of secondary antibodies tagged with **3-NHS** and the NHS ester of a highly photostable anionic rhodamine dye *abberior STAR 512*,^19^ respectively, with the actin cytoskeleton and nuclei counterstained with *SiR-actin*^20^ and the DNA binder 4′,6-diamidino-2-phenylindole (DAPI). Two-color imaging could be performed first, followed by photoactivation of **3** with 405 nm laser line and recording the data for **3** and DAPI (Figure S9a). In living U2OS-Vim-Halo cells, photoactivatable HaloTag ligand **3-Halo** was employed along with live-cell compatible pentamethine cyanine mitochondrial label *MitoTracker Deep Red FM*,^21^ a cell-permeant nuclear counterstain Hoechst 33342 and the non-covalent small molecule *abberior LIVE 510-tubulin* probe (a cabazitaxel-tagged rhodamine dye^22^). Since the latter is less photostable as compared to *STAR 512*, partial bleaching of the labeled vimentin filaments was observed in the second acquisition (Figure S9b). In both cases, complete color separation in all four channels required the use of spectral unmixing.^12^

**Figure 4.**
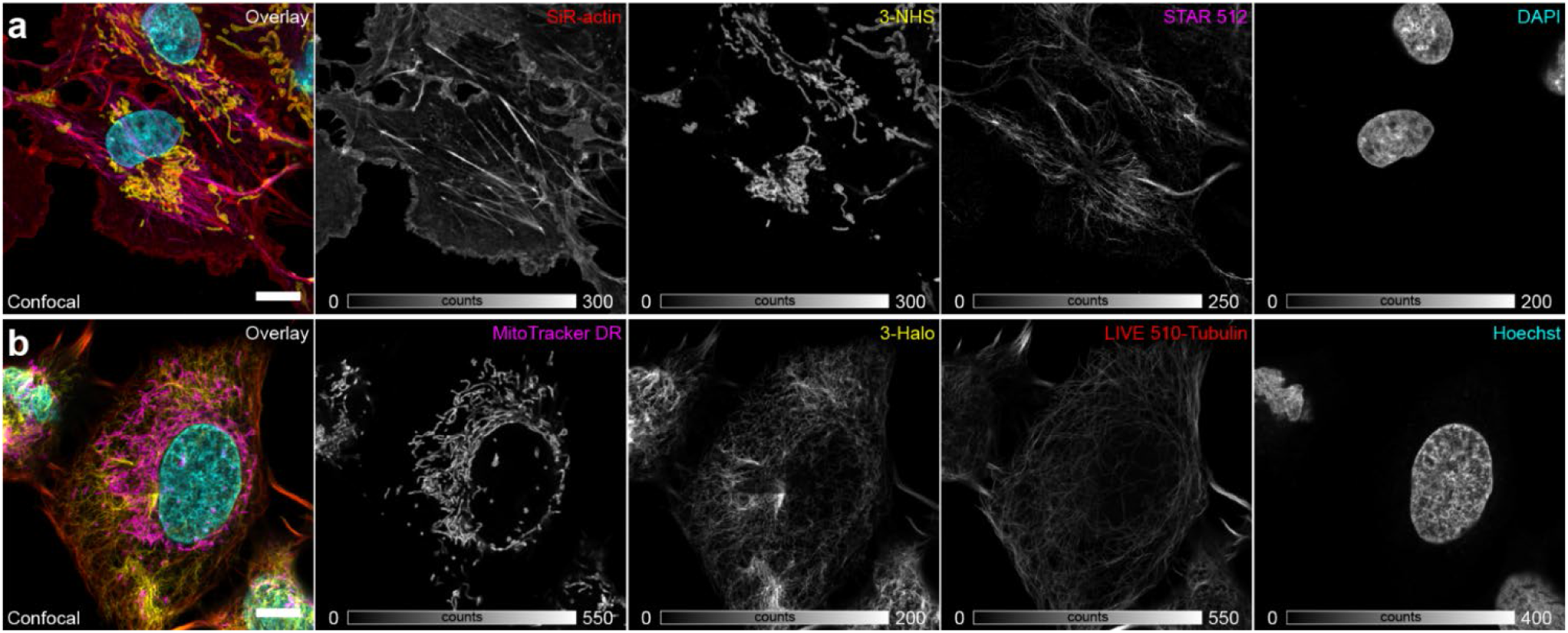
Multicolor imaging with photoactivatable large Stokes shift labels. (a) Confocal imaging of fixed COS-7 cells labeled by indirect immunofluorescence for mitochondria (**3-NHS**) and vimentin (*abberior STAR 512* NHS ester), and with small-molecule fluorescent probes *SiR-actin* (for F-actin) and DAPI (for nuclear DNA). (b) Confocal imaging of living U2OS cells stably expressing a vimentin-HaloTag construct labeled with **3-Halo** and live cell-compatible probes *MitoTracker Deep Red FM* (specific for mitochondria), *abberior LIVE 510-tubulin* (microtubules) and Hoechst 33342 (DNA). Spectral unmixing of individual channels was performed according to reference ^12^. Scale bars: 10 µm.

In conclusion, we introduced live-cell membrane-permeant photoactivatable fluorescent dyes with large Stokes shifts (∼100 nm) based on a caging group-free activation mechanism unique for PaX fluorophores.^8^ Similarly to these small Stokes shift fluorophores (<40 nm), the best-performing dye **3** was found compatible with both STED (at 775 nm) and PALM fluorescence nanoscopy, each imposing distinct demands on the fluorescent label. Photoactivatable ligands derived from dye **3** permit up to four-color microscopy in fixed and live samples and form two-color combinations with less photostable commercial large Stokes shift probes based on a photobleaching-photoactivation sequence, which can be trivially realized with commercial fluorescence microscopes. We also anticipate the application of our photoactivatable large Stokes shift labels in multicolor single-molecule tracking of labeled biomolecules, further expanding the toolkit available for studying biochemical mechanisms.

## ASSOCIATED CONTENT

**Supporting Information**. Additional experimental details, materials, methods, and characterization data for all new compounds (PDF).

## Notes

The authors declare the following competing financial interest(s): R.L., M.L.B. and A.N.B. are co-inventors of a patent application covering the photoactivatable dyes of this work, filed by the Max Planck Society. S.W.H. owns shares of Abberior GmbH and Abberior Instruments GmbH, whose dyes and STED microscope, respectively, have been used in this study. The remaining authors declare no competing interests.

## ACKNOWLEDGMENT

The research was funded by Bundesministerium für Bildung und Forschung (German Federal Ministry of Education and Research), project no. 13N14122 ‘3D Nano Life Cell’ (to S.W.H.). R.L. is grateful to the Max Planck Society for a Nobel Laureate Fellowship. We thank Prof. S. Jakobs (Max Planck Institute for Multidisciplinary Sciences, University of Göttingen) for providing the U2OS-Vim-Halo and U2OS-Vim-SNAP cells and the European Molecular Biology Laboratory for providing the U2OS-NUP96-Halo cells. We thank the staff of Mass Spectrometry Core Facility (Max Planck Institute for Medical Research) for recording mass spectra of small molecules, the Optical Microscopy Facility (MPI MR) for the use of their fluorescence microscopes and M. Remmel (MPI MR) for the use of a custom-built PALM microscope.

## REFERENCES

1 >Sednev, M. V.; Belov, V. N.; Hell, S. W. Fluorescent dyes with large Stokes shifts for super-resolution optical microscopy of biological objects: a review. Methods Appl. Fluoresc. 2015, 3, 042004.

2 Croce, A.C.; Bottiroli, G. Autofluorescence spectroscopy and imaging: biomedical research and diagnosis. Eur. J. Histochem. 2014, 58, 320–337.

3 Chen, H.; Liu, L.; Qian, k.; Liu, H.; Wang, Z.; Gao, F.; Qu, C.; Dai, W.; Lin, D.; Chen, K.; Liu, H.; Cheng, Z. Bioinspired large Stokes shift small molecular dyes for biomedical fluorescence imaging. Sci. Adv. 2022, 8, eabo3289.

4 Dhara, A.; Sadhukhan, T.; Sheetz, E. G.; Olsson, A. H.; Raghavachari, K.; Flood, A. H. Zero-overlap fluorophores for fluorescent studies at any concentration. J. Am. Chem. Soc. 2020, 142, 12167−12180.

5 (a)Chozinski, T. J.; Gagnon, L. A.; Vaughan, J. C. Twinkle, twinkle little star: Photoswitchable fluorophores for super-resolution imaging. FEBS Letters 2014, 588, 3606–3612. (b)Zhang, Y.; Raymo, F. M. Photoactivatable fluorophores for single-molecule localization microscopy of live cells. Methods Appl. Fluoresc. 2020, 8, 032002.

6 Klötzner, D.-P.; Klehs, K.; Heilemann, M.; Heckel, A. A new photoactivatable near-infrared-emitting QCy7 fluorophore for single-molecule super-resolution microscopy. Chem. Commun. 2017, 53, 9874– 9877.

7 Tran, M. N.; Rarig, R.-A. F.; Chenoweth, D. M. Synthesis and properties of lysosome-specific photoactivatable probes for live-cell imaging. Chem. Sci. 2015, 6, 4508–4512.

8 Lincoln, R.; Bossi, M. L.; Remmel, M.; D’Este, E.; Butkevich, A. N.; Hell, S. W. A general design of caging-group-free photoactivatable fluorophores for live-cell nanoscopy. Nat. Chem. 2022, 14, 1013– 1020.

9 Butkevich, A. N.; Lukinavičius, G.; D’Este, E.; Hell, S. W. Cell-permeant large Stokes shift dyes for transfection-free multicolor nanoscopy. J. Am. Chem. Soc. 2017, 139, 12378−12381.

10 Horváth, P.; Šebej, P.; Šolomek, T.; Klán, P. Small-molecule fluorophores with large Stokes shifts: 9-iminopyronin analogues as clickable tags. J. Org. Chem. 2015, 80, 1299–1311.

11 Wu, L.; Burgess, K. Fluorescent Amino- and Thiopyronin Dyes. Org. Lett. 2008, 10, 1779–1782.

12 Walter, J. Spectral Unmixing for ImageJ – documentation (v1.2), https://imagej.nih.gov/ij/plugins/docs/SpectralUnmixing.pdf (accessed November 2022).

13 Thevathasan, J. V.; Kahnwald, M.; Cieslinski, K.; Hoess, P.; Peneti, S. K.; Reitberger, M.; Heid, D.; Kasuba, K. C.; Hoerner, S. J.; Li, Y. M.; Wu, Y. L.; Mund, M.; Matti, U.; Pereira, P. M.; Henriques, R.; Nijmeijer, B.; Kueblbeck, M.; Jimenez Sabinina, V.; Ellenberg, J.; Ries, J. Nuclear pores as versatile reference standards for quantitative superresolution microscopy. Nat. Methods, 2019, 16, 1045–1053.

14 Los, G. V.; Encell, L. P.; McDougall, M. G.; Hartzell, D. D.; Karassina, N.; Zimprich, C.; Wood, M. G.; Learish, R.; Ohana, R. F.; Urh, M.; Simpson, D.; Mendez, J.; Zimmerman, K.; Otto, P.; Vidugiris, G.; Zhu, J.; Darzins, A.; Klaubert, D. H.; Bulleit, R. F.; Wood, K. V. HaloTag: A Novel Protein Labeling Technology for Cell Imaging and Protein Analysis. ACS Chem. Biol. 2008, 3, 373–382.

15 Butkevich, A. N.; Ta, H.; Ratz, M.; Stoldt, S.; Jakobs, S.; Belov, V. N.; Hell, S. W. Two-color 810 nm STED nanoscopy of living cells with endogenous SNAP-tagged fusion proteins. ACS Chem. Biol. 2018, 13, 475–480.

16 Gautier, A.; Juillerat, A.; Heinis, C.; Corrêa Jr., I. R.; Kindermann, M.; Beaufils, F.; Johnsson, K. An engineered protein tag for multiprotein labeling in living cells. Chem. Biol. 2008, 15, 128–136.

17 Marciniszyn Jr., J.; Hartsuck, J. A.; Tang, J. Mode of inhibition of acid proteases by pepstatin. J. Biol. Chem. 1976, 251, 7088–7094.

18 Lukinavičius, G.; Reymond, L.; Umezawa, K.; Sallin, O.; D’Este, E.; Göttfert, F.; Ta, H.; Hell, S. W.; Urano, Y.; Johnsson, K. Fluorogenic probes for multicolor imaging in living cells. J. Am. Chem. Soc. 2016, 138, 9365–9368.

19 Mitronova, G.Y.; Belov, V.N.; Bossi, M.L.; Wurm, C.A.; Meyer, L.; Medda, R.; Moneron, G.; Bretschneider, S.; Eggeling, C.; Jakobs, S.; Hell, S.W. New fluorinated rhodamines for optical microscopy and nanoscopy. Chem. Eur. J. 2010, 16, 4477–4488.

20 Lukinavičius, G.; Reymond, L.; D’Este, E.; Masharina, A.; Göttfert, F.; Ta, H.; Güther, A.; Fournier, M.; Rizzo, S.; Waldmann, H.; Blaukopf, C.; Sommer, C.; Gerlich, D. W.; Arndt, H.-D.; Hell, S. W.; Urano, Y.; Johnsson, K. Fluorogenic probes for live-cell imaging of the cytoskeleton. Nat. Methods 2014, 11, 731–733.

21 Shim, S.-H.; Xia, C.; Zhong, G.; Babcock, H. P.; Vaughan, J. C.; Huang, B.; Wang, X.; Xu, C.; Bi, G.-Q.; Zhuang, X. Super-resolution fluorescence imaging of organelles in live cells with photoswitchable membrane probes. Proc. Natl. Acad. Sci. USA, 2012, 109, 13978–13983.

22 Grimm, F.; Nizamov, S.; Belov, V. N. Green-emitting rhodamine dyes for vital labeling of cell organelles using STED super-resolution microscopy. ChemBioChem 2019, 20, 2248–2254.

