## Supplementary Material for "Photoactivatable Large Stokes Shift Fluorophores for Multicolor Nanoscopy"

### Photoactivatable large Stokes shift fluorescent dyes for multicolor nanoscopy

#### Supplementary Figures

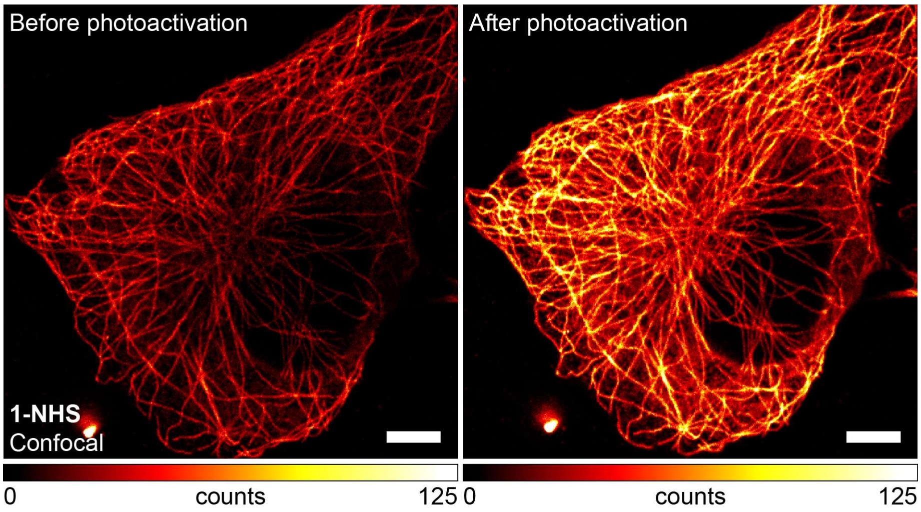

**Supplementary Figure S1.** Confocal imaging with large Stokes shift dye **1** showing high initial fluorescence (before activation). Confocal images of tubulin filaments in methanol-fixed COS-7 cells labelled by indirect immunofluorescence with antibody conjugates of compound **1-NHS** imaged before (left) and after (right) photoactivation ($\lambda_{act}$ = 405 nm). Scale bars: 5 µm.

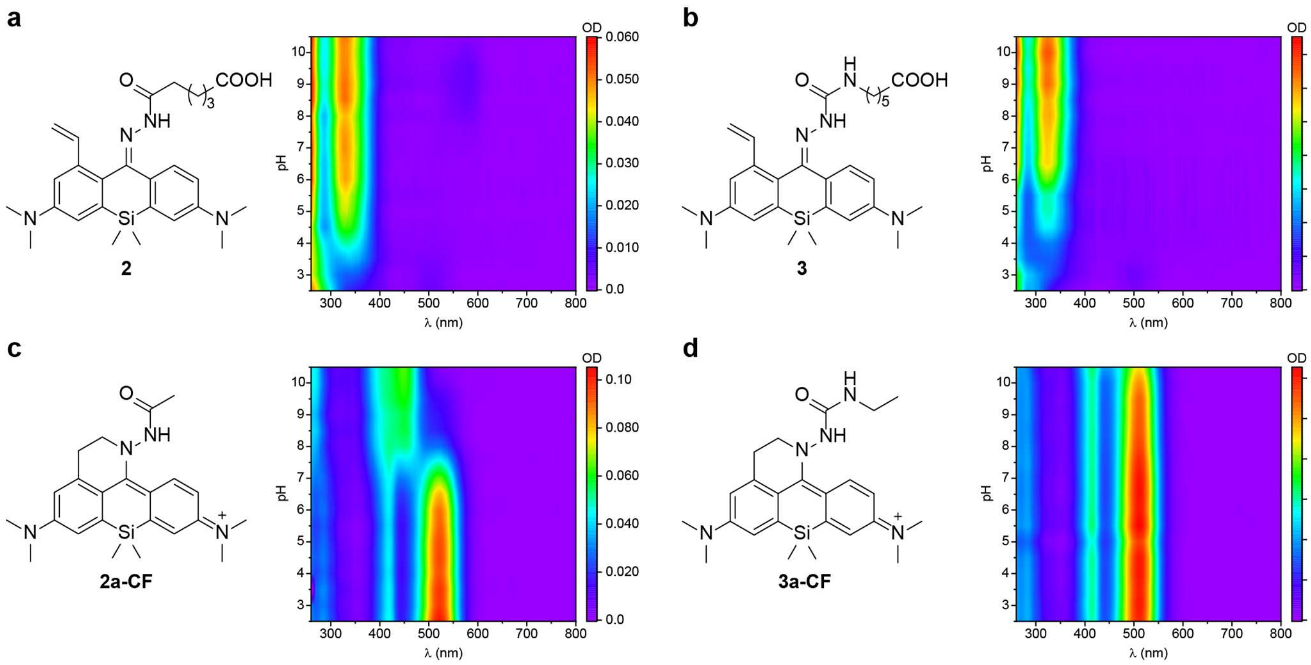

**Supplementary Figure S2.** Absorption spectral changes of 5.0 µM solutions of **2** (a) and **3** (b) and their corresponding representative closed form analogues **2a-CF** (c) and **3a-CF** (d) measured at different pH values in phosphate buffered saline containing 10% DMSO (v/v).

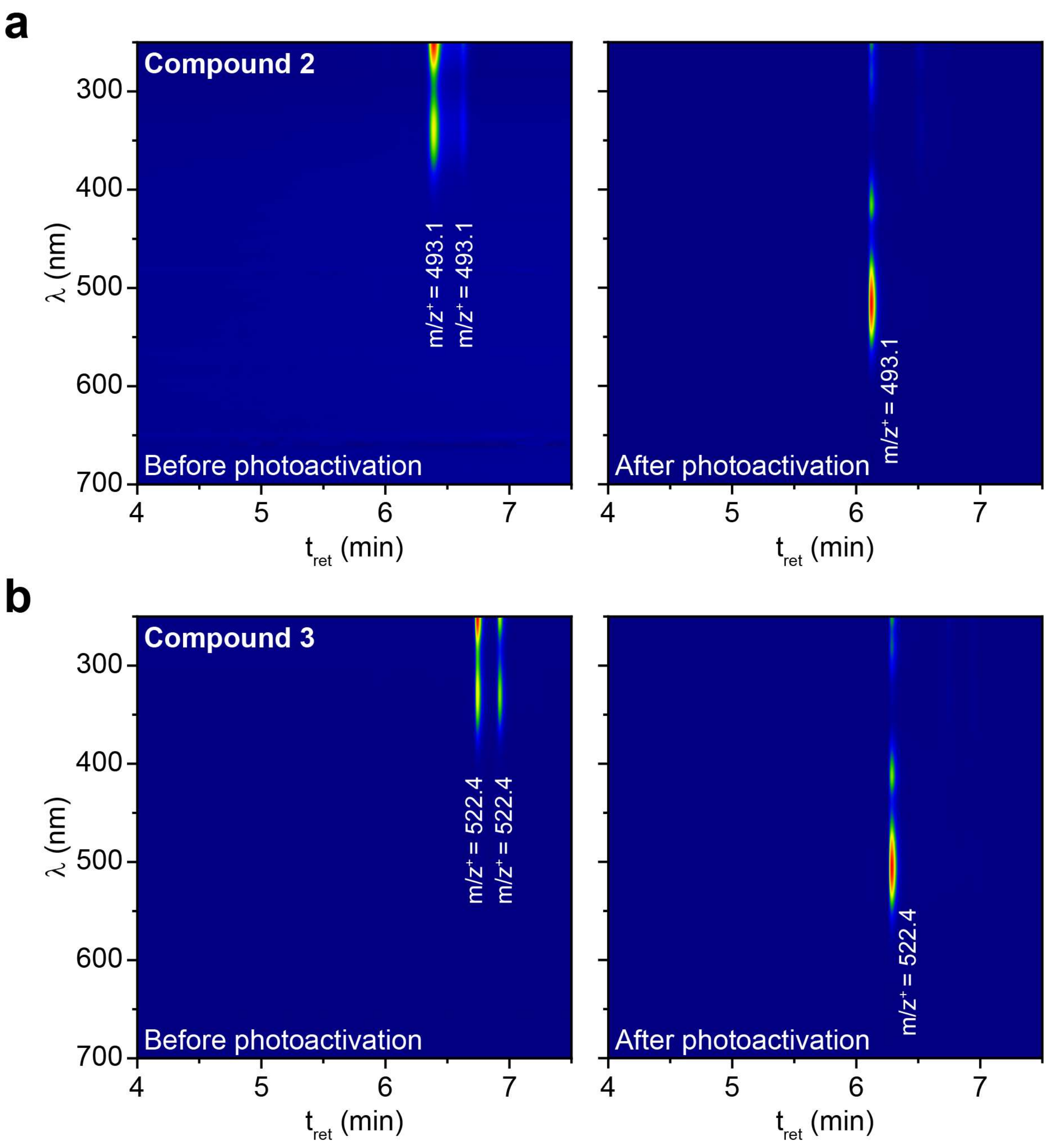

**Supplementary Figure S3.** HPLC-MS chromatograms of solutions of **2** and **3** (6.7 µM) in phosphate buffer (100 mM, pH 7) before and after irradiation ($\lambda_{act}$ = 365 nm). The double peaks of **2** and **3** before activation correspond to the (*E*/*Z*)-isomers of the hydrazones **2-OF**, **3-OF**.

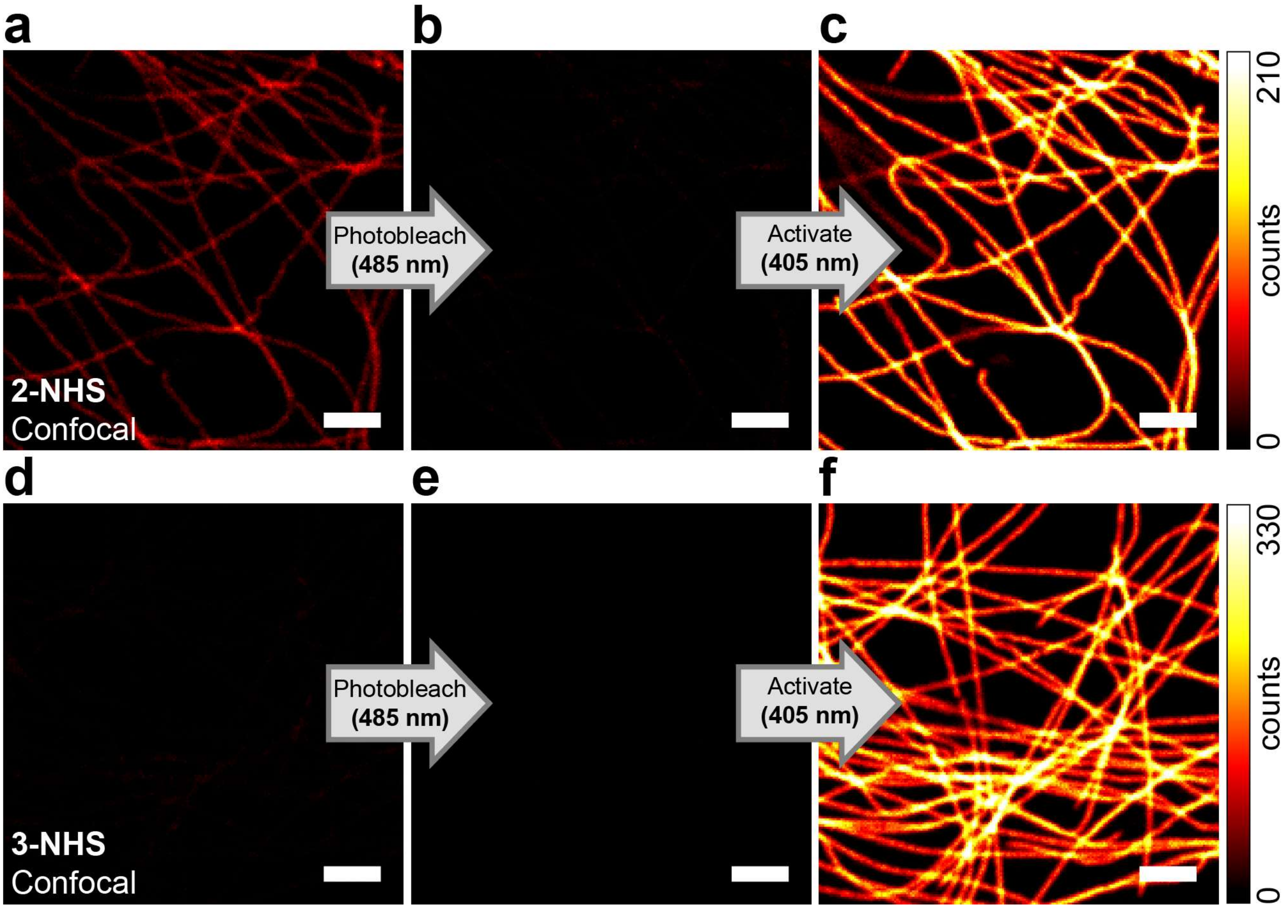

**Supplementary Figure S4.** One-photon activation of large Stokes shift dyes **2** and **3**. Confocal images of tubulin filaments in fixed COS-7 cells labelled by indirect immunofluorescence with antibody conjugates of compound **2-NHS** (a-c) and **3-NHS** (d-f) imaged before (a, d) and after (b, e) photobleaching ($\lambda_{excit}$ = 485 nm), and subsequent photoactivation (c, f; $\lambda_{act}$ = 405 nm). Scale bars: 2 µm.

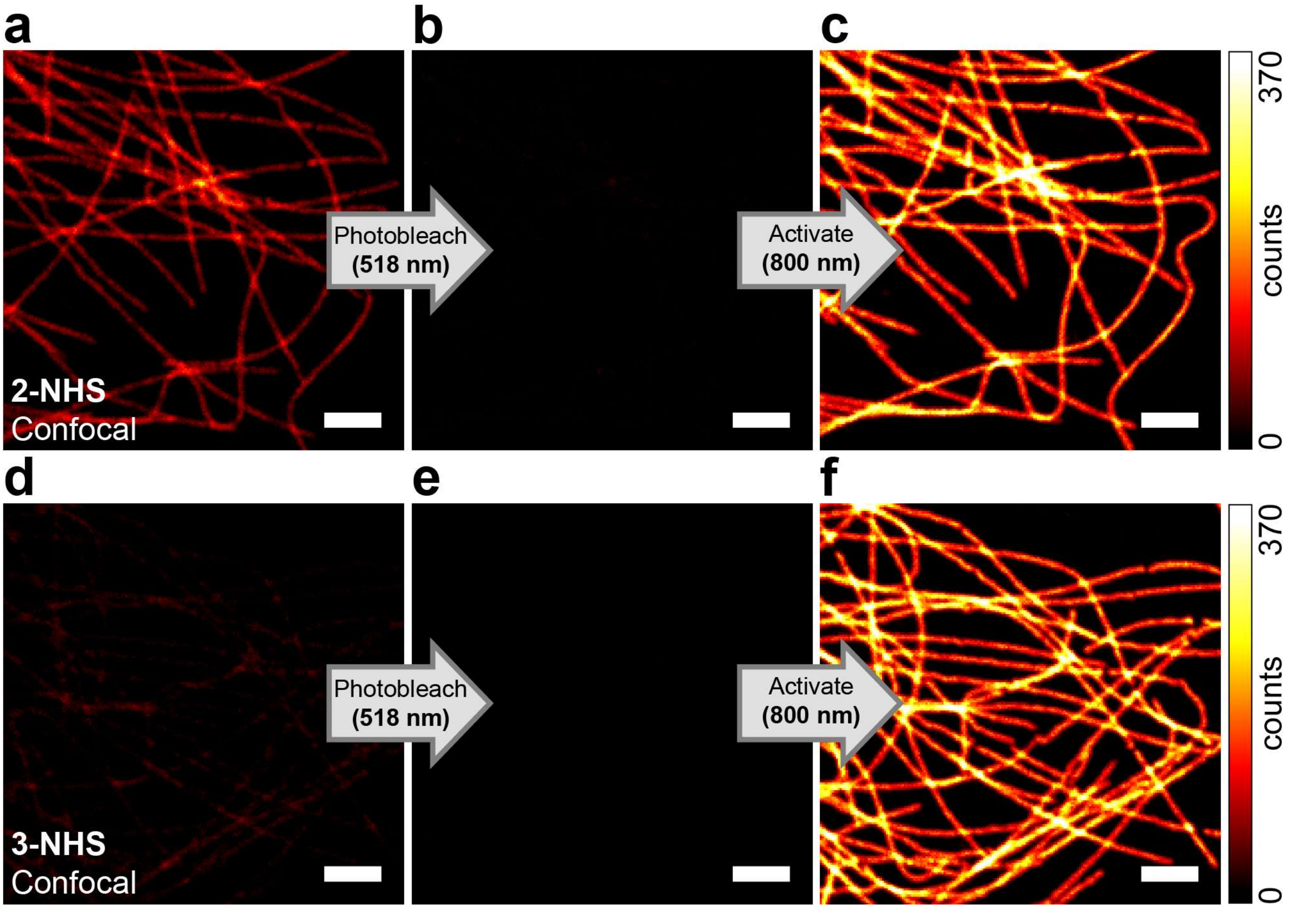

**Supplementary Figure S5.** Two-photon activation of large Stokes shift dyes **2** and **3**. Confocal images of tubulin filaments in fixed COS-7 cells labelled by indirect immunofluorescence with antibody conjugates of compound **2-NHS** (a-c) and **3-NHS** (d-f) imaged before (a, d) and after (b, e) photobleaching ($\lambda_{excit}$ = 518 nm), and subsequent photoactivation (c, f; $\lambda_{act}$ = 800 nm). Scale bars: 2 μm.

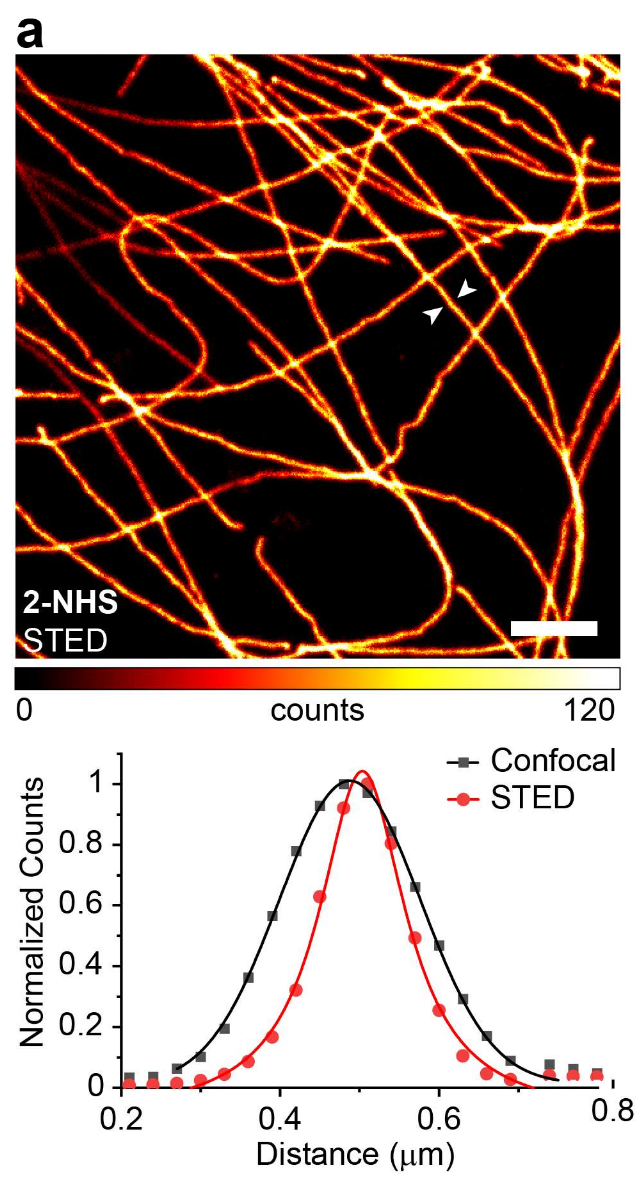

**Supplementary Figure S6.** STED images of tubulin filaments in fixed COS-7 cells labeled by indirect immunofluorescence with secondary antibodies tagged with **2-NHS** (a) or **3-NHS** (b) and corresponding intensity profiles across selected regions of the images (marked with arrows) fit to a Gaussian and Lorentzian model for the confocal and STED data, respectively .

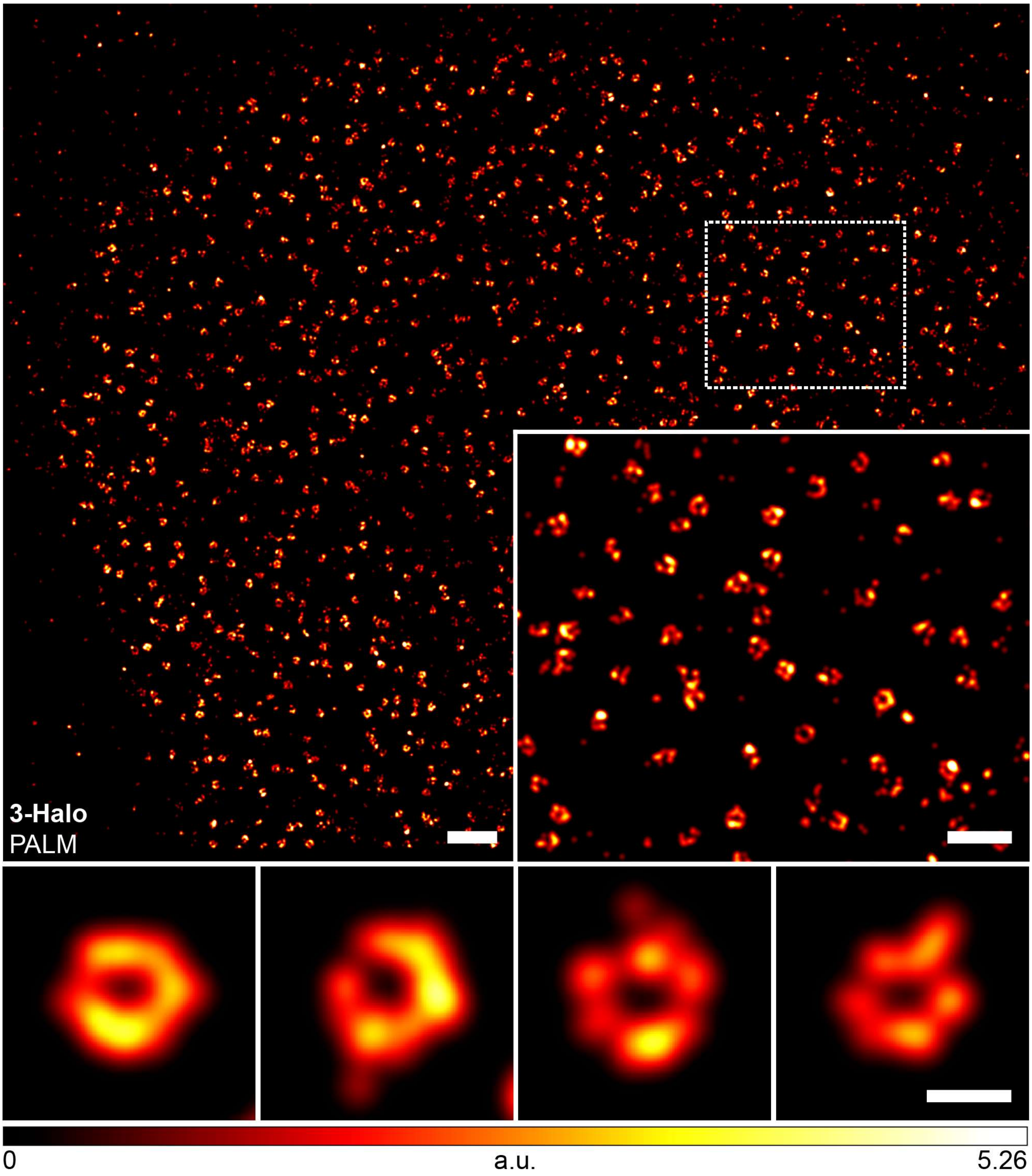

**Supplementary Figure S7.** PALM imaging of nuclear pore complexes (NPCs) in fixed cells using photoactivatable large Stokes shift HaloTag label **3-Halo**. PALM image of fixed U2OS cells stably expressing a NUP96-HaloTag construct labeled with **3-Halo** (200 nM, overnight). Inset: Magnified region marked in overview image. Bottom row: Magnified individual NPCs. Scale bars: 1 μm (overview), 500 nm (inset), 100 nm (bottom row).

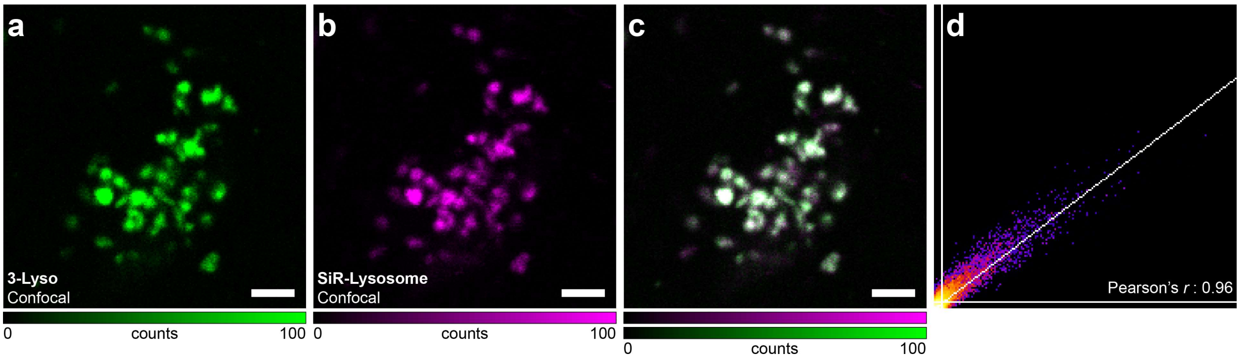

**Supplementary Figure S8.** Confocal images of living U2OS cells co-stained with **3-Lyso** (a, 20 nM overnight) and SiR-Lysosome (b, 500 nM for 60 min), overlay (c) and corresponding Pearson correlation analysis (d). Scale bars: 2 µm.

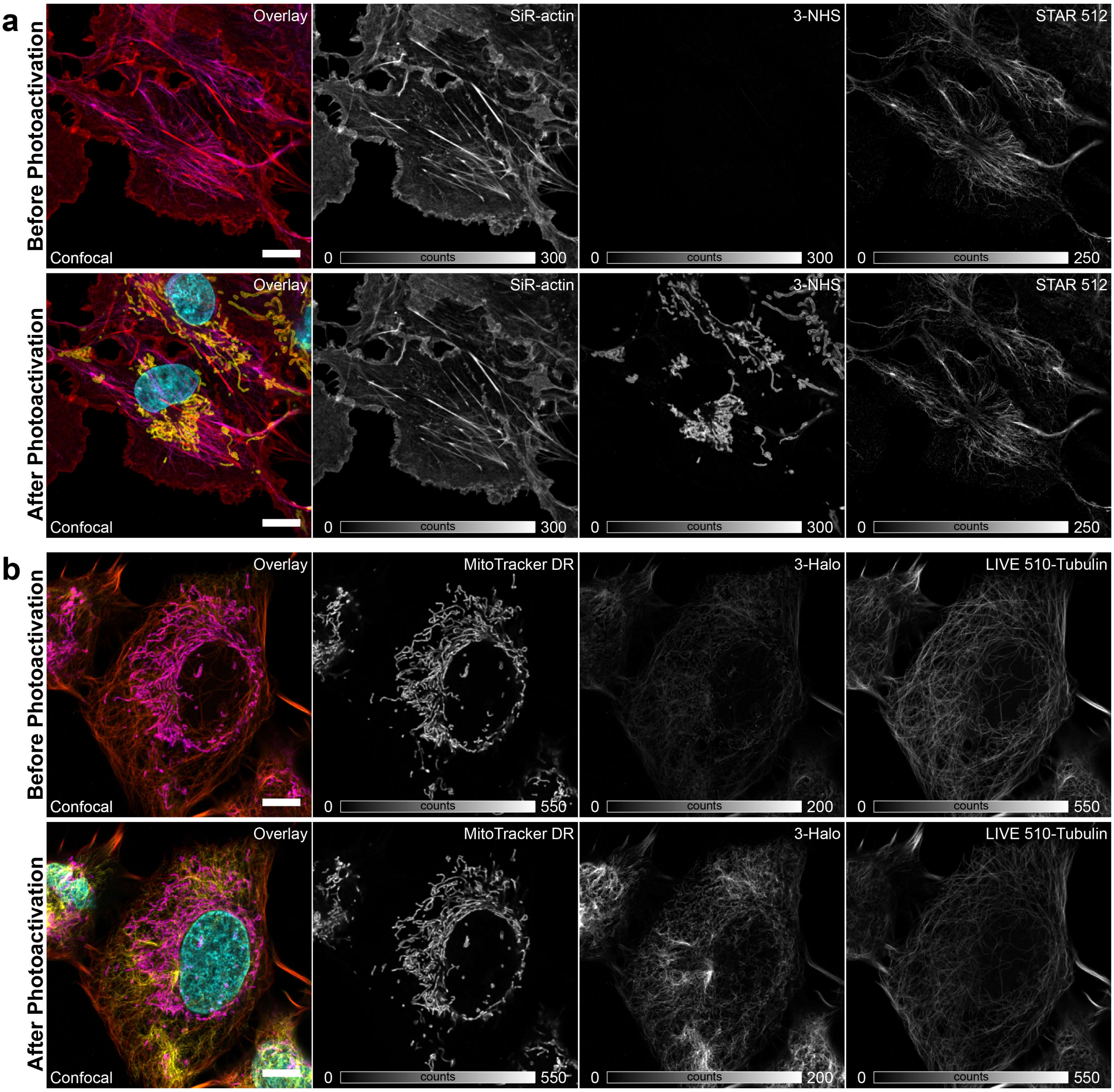

**Supplementary Figure S9.** Effect of photoactivation of large Stokes shift labels on commercial fluorophores during multicolor imaging. (a) Confocal imaging of fixed COS-7 cells labeled by indirect immunofluorescence for mitochondria (**3-NHS**) and vimentin (*abberior STAR 512*), and small-molecule dyes for actin (*SiR-actin*) and nuclei (DAPI) before (top row) and after (bottom row) photoactivation of **3** ($\lambda_{act}$ = 405 nm); DAPI channel (excitation 405 nm) was imaged only after photoactivation (to avoid the inadvertent photoactivation of **3**). (b) Confocal imaging of U2OS cells stably expressing a vimentin-HaloTag construct labelled with live-cell dyes for mitochondria (MitoTracker Deep Red FM), vimentin (**3-Halo**), microtubules (*abberior LIVE 510 tubulin*), before (top row) and after (bottom row) photoactivation of **3** ($\lambda_{act}$ = 405 nm); Hoechst channel was imaged only after photoactivation. Spectral unmixing of individual channels was performed according to [1]. Scale bars: 10 μm.

#### Supplementary Tables

##### Supplementary Table 1. Confocal and STED imaging parameters.

| Figure | Micro-scope | Compound | Photoactivation | Excitation | | Depletion | | Detection windows (nm) | Pixel size (nm) | Dwell time (μs) | Line reps. |
|---|---|---|---|---|---|---|---|---|---|---|---|
| | | | | Laser (nm) | Power (μW) | Laser (nm) | Power (mW) | | | | |
| Fig. 2b | 1 | **2-NHS** (Confocal)<br>**2-NHS** (STED) | 405 nm (13 μW) | 485<br>485 | 6.37<br>6.37 | -<br>775 | -<br>150 | 560–700<br>560–700 | 70<br>30 | 20<br>20 | 2x1<br>2x3 |
| Fig. 2c | 1 | **3-NHS** (Confocal)<br>**3-NHS** (STED) | 405 nm (13 μW) | 485<br>485 | 6.37<br>6.37 | -<br>775 | -<br>230 | 560–700<br>560–700 | 70<br>30 | 20<br>20 | 2x1<br>2x3 |
| Fig. 2d | 1 | **3-Halo** (Confocal) | 405 nm (13 μW) | 485 | 20.0 | - | - | 560–700 | 70 | 40 | 4x1 |
| Fig. 2e | 1 | **3-BG** (Confocal) | 405 nm (13 μW) | 485 | 20.0 | - | - | 560–700 | 70 | 40 | 4x1 |
| Fig. 2f | 1 | **3-Tz** (Confocal) | 405 nm (13 μW) | 485 | 20.0 | - | - | 560–700 | 70 | 40 | 4x1 |
| Fig. 3 | 1 | DyLight 515-LS<br>**3-NHS** (Confocal)<br>**3-NHS** (STED) | -<br>405 nm (13 μW) | 485<br>485<br>485 | 6.37<br>6.37<br>6.37 | -<br>-<br>775 | -<br>-<br>230 | 560–700<br>560–700<br>560–700 | 70<br>70<br>30 | 20<br>20<br>20 | 2x1<br>2x1<br>2x3 |
| Fig. 4a | 1 | SiR-Actin<br>**3-NHS**<br>STAR 512<br>DAPI | -<br>405 nm (13 μW)<br>-<br>- | 640<br>485<br>485<br>405 | 0.26<br>13.2<br>6.37<br>209 | -<br>-<br>-<br>- | -<br>-<br>-<br>- | 650–750<br>600–700<br>500–600<br>415–500 | 70<br>70<br>70<br>70 | 20<br>20<br>20<br>20 | 2x1<br>2x1<br>2x1<br>2x1 |
| Fig. 4b | 1 | MitoTracker DR<br>**3-Halo**<br>Live 510-Tubulin<br>Hoechst | -<br>405 nm (13 μW)<br>-<br>- | 640<br>485<br>485<br>405 | 1.32<br>20.0<br>6.37<br>209 | -<br>-<br>-<br>- | -<br>-<br>-<br>- | 650–750<br>590–700<br>500–590<br>415–500 | 70<br>70<br>70<br>70 | 20<br>20<br>20<br>20 | 2x1<br>2x1<br>2x1<br>2x1 |

| | | | | | | | | | | | |
|---|---|---|---|---|---|---|---|---|---|---|---|
| Fig. S1 | 1 | **1-NHS** | 405 nm (13 μW) | 485 | 16.5 | - | - | 520–700 | 70 | 20 | 2x1 |
| Fig. S4a-c | 1 | **2-NHS** (Confocal) | 405 nm (13 μW) | 485 | 6.37 | - | - | 560–700 | 70 | 20 | 2x1 |
| Fig. S4d-f | 1 | **3-NHS** (Confocal) | 405 nm (13 μW) | 485 | 6.37 | - | - | 560–700 | 70 | 20 | 2x1 |
| Fig. S5a-c | 2 | **2-NHS** (Confocal) | 800 nm (44.6 mW) | 518 | 1.6 | - | - | 560–700 | 70 | 20 | 2x1 |
| Fig. S5d-f | 2 | **3-NHS** (Confocal) | 800 nm (44.6 mW) | 518 | 1.6 | - | - | 560–700 | 70 | 20 | 2x1 |
| Fig. S6a | 1 | **2-NHS** (STED) | 405 nm (13 μW) | 485 | 6.37 | 775 | 150 | 560–700 | 30 | 20 | 2x3 |
| Fig. S6b | 1 | **3-NHS** (STED) | 405 nm (13 μW) | 485 | 6.37 | 775 | 230 | 560–700 | 30 | 20 | 2x3 |
| Fig. S8 | 1 | **3-Lyso** | 405 nm (13 μW) | 485 | 6.37 | - | - | 560–650 | 70 | 20 | 2x1 |
| | | SiR-Lysosome | - | 640 | 32.8 | - | - | 650–750 | 70 | 20 | 2x1 |
| Fig. S9a | 1 | SiR-Actin | - | 640 | 0.26 | - | - | 650–750 | 70 | 20 | 2x1 |
| | | **3-NHS** | 405 nm (13 μW) | 485 | 13.2 | - | - | 600–700 | 70 | 20 | 2x1 |
| | | STAR 512 | - | 485 | 6.37 | - | - | 500–600 | 70 | 20 | 2x1 |
| | | DAPI | - | 405 | 209 | - | - | 415–500 | 70 | 20 | 2x1 |
| Fig. S9b | 1 | MitoTracker DR | - | 640 | 1.32 | - | - | 650–750 | 70 | 20 | 2x1 |
| | | **3-Halo** | 405 nm (13 μW) | 485 | 20.0 | - | - | 590–700 | 70 | 20 | 2x1 |
| | | Live 510-Tubulin | - | 485 | 6.37 | - | - | 500–590 | 70 | 20 | 2x1 |
| | | Hoechst | - | 405 | 209 | - | - | 415–500 | 70 | 20 | 2x1 |

#### Supplementary Table 2. Antibodies and nanobodies used.

| Reagent | Type | Target | Host | Supplier | Catalogue No. | Dilution |
|---|---|---|---|---|---|---|
| **AffiniPure Goat Anti-Rabbit IgG (H+L)** | Secondary antibody | Rabbit | Goat | Jackson ImmunoResearch Europe Ltd. | 111-005-003 | 1:100 to 1:500 |
| **AffiniPure Goat Anti-Mouse IgG (H+L)** | Secondary antibody | Mouse | Goat | Jackson ImmunoResearch Europe Ltd. | 115-005-003 | 1:100 to 1:500 |
| **Anti-alpha Tubulin antibody** | Primary Antibody (polyclonal) | α-tubulin | Rabbit | Abcam | ab18251 | 1:200 |
| **Anti-TOMM20 antibody** | Primary Antibody (monoclonal) | TOMM20 | Rabbit | Abcam | ab186735 | 1:200 |
| **Anti-alpha-Tubulin antibody** | Primary Antibody (monoclonal) | α-tubulin | Mouse | Synaptic Systems | 302 211 | 1:200 |
| **Anti-Vimentin antibody** | Primary Antibody (monoclonal) | Vimentin | Mouse | Sigma-Aldrich | V6389 | 1:200 |
| **Anti-mouse STAR 512** | Secondary antibody | Mouse | Goat | Abberior GmbH | 2-0012-001-0 | 1:500 |

#### Supplementary Methods

##### General experimental information and synthesis

**NMR spectra** were recorded at 25 °C with a Bruker DPX 400 spectrometer using the Bruker Topspin 3.5 software at 400.15 MHz ($^{1}H$), 376.48 MHz ($^{19}F$) and 100.63 MHz ($^{13}C$) and are reported in ppm. All $^{1}H$ spectra are referenced to tetramethylsilane ($\delta$ = 0 ppm) added as an internal standard or using the signals of the residual protons of $CHCl_3$ (7.26 ppm) in $CDCl_3$, $CHD_2CN$ (1.94 ppm) in $CD_3CN$ or DMSO-$d_5$ (2.50 ppm) in DMSO-$d_6$. $^{13}C$ spectra are referenced to tetramethylsilane ($\delta$ = 0 ppm) using the signals of the solvent: $CDCl_3$ (77.16 ppm), $\underline{C}D_3CN$ (1.32 ppm) or DMSO-$d_6$ (39.52 ppm). Multiplicities of signals are described as follows: s = singlet, d = doublet, t = triplet, q = quartet, p = pentet, m = multiplet or overlap of non-equivalent resonances; br = broad signal. Coupling constants ($J$) are given in Hz.

**ESI-MS** (low resolution mass spectra with electrospray ionization) were recorded on a Shimadzu LC-MS system described below. **ESI-HRMS** (high resolution mass spectra) were recorded on a maXis II ETD instrument (Bruker) at the Mass Spectrometry Core facility of the Max Planck Institute for Medical Research (Heidelberg, Germany).

**Liquid chromatography:** Analytical liquid chromatography-mass spectrometry was performed on an LC-MS system (Shimadzu, controlled with LabSolutions 5.89 software): 2x LC-20AD HPLC pumps with DGU-20A3R solvent degassing unit, SIL-20ACHT autosampler, CTO-20AC column oven, SPD-M30A diode array detector and CBM-20A communication bus module, integrated with CAMAG TLC-MS interface 2, FCV-20$AH_2$ diverter valve and LCMS-2020 spectrometer with electrospray ionization (ESI, 100 – 1500 $m/z$). Analytical column: Hypersil GOLD 50×2.1 mm 1.9µm, standard conditions: sample volume 1-2 µL, solvent flow rate 0.5 mL/min, column temperature 30 °C. General method: isocratic 95:5 A:B over 2 min, then gradient 95:5 – 0:100 A:B over 5 min, then isocratic 0:100 A:B over 2 min; solvent A = water + 0.1% v/v $HCO_2H$, solvent B = acetonitrile + 0.1% v/v $HCO_2H$.

Preparative HPLC was performed on an Interchim puriFlash 5.250P preparative HPLC/Flash hybrid system (Article No. PFG5Q0, Interchim) with a 5 mL injection loop, a 200-800 nm UV-Vis detector and an integrated ELSD detector (Article No. ELSD01, Interchim), using the preparative column Interchim 250×21.2 mm 5 µm Uptisphere

Strategy PhC4 (Article No. US5PHC4-250/212, Interchim), unless specified otherwise, and the conditions as indicated for individual preparations.

**Analytical TLC** was performed on Merck Millipore ready-to-use plates with silica gel 60 ($F_{254}$) (Cat. No. 1.05554.0001).

**Preparative flash chromatography** was performed on Biotage Isolera Spektra One flash purification system (Biotage AG, Sweden) using the Puriflash Silica HP 30µm series flash cartridges from Interchim and solvent gradients as indicated.

For **photoactivation** of the dyes on a semipreparative scale, Photoreactor m1 (Penn PhD, USA) was used together with a custom home-built 405 nm LED unit (described in [2]).

**Warning:** All of the following photoactivatable dyes and intermediates are light sensitive, in particular against ultraviolet and blue light. The exposure of solutions to direct or reflected sunlight or the light of luminescent lamps resulted in inadvertent photoactivation. All reactions were performed in amber glassware or shielded from the daylight with aluminum foil, and generic 12 V red LED strips (IP65 waterproof, 620-640 nm) were used for ambient lighting of the lab. Similar precautions should be taken when labeling, transferring the label solutions or samples labelled with the photoactivatable dyes of the present study.

##### Optical spectroscopy

Fluorescence quantum yield measurements were measured Quantaurus-QY absolute PL quantum yield spectrometer (C11347-11, Hamamatsu) in methanol and phosphate buffer (pH = 7). Fluorescence lifetimes were measured with a FluoTime 300 fluorescence lifetime spectrometer (PicoQuant, controlled with the EasyTau1.4 software) in 3 mL quartz cells (optical path length 1 cm, model 119F-10-40 or model 111-10-40, Hellma Analytics). All measurements were performed in air-saturated solvents at ambient temperature.

#### Photolysis of compounds and chemometric analysis of the photoactivation kinetics

Solutions in phosphate buffer (100 mM, pH = 7.0; 6.7 µM dye) or methanol were irradiated in a previously described [3] home-built setup with a 405 nm LED source (M405L3, Thorlabs Inc.) in combination with a bandpass (10 nm) filter (FB405-10, Thorlabs Inc.). During the irradiation, samples were maintained at 20 °C and continuously stirred with a Peltier-based temperature-controlled cuvette holder (Luma 40, Quantum Northwest, Inc.). The absorption and emission of irradiated solutions was monitored at desired irradiation intervals with a fiber-based spectrometer (Flame-S-UV-Vis-ES, Ocean Insight). For absorption measurements, a deuterium and tungsten halogen source was used for illumination (DH-2000-BAL, Ocean Insight), and for fluorescence excitation was performed in a 90° configuration with an LED source (Thorlabs Inc.) in combination with an appropriate bandpass (10 nm) filter emitting at a wavelength suitable for each compound. Data collection and analysis was performed with custom-made routines in Matlab. Samples for LCMS or ESI-MS analysis were taken before and after photolysis was performed.

#### Determination of pH-dependent behavior by absorption spectroscopy

A series of buffers ranging from pH 2.5 to 10.5 was prepared from PBS adjusted with HCl or NaOH (3 M and 0.3 M) and monitoring with a PT-10 pH-meter (Sartorius AG, Göttingen, Germany). Using these buffers, 5 µM solutions of the dyes (**2**, **3**, **2-CF** or **3-CF**) were prepared in 96-well UV-STAR® COC microplate plates with flat-bottoms (Greiner Bio-One GmbH, Frickenhausen, Germany) containing 10% (v/v) DMSO. Each pH condition was prepared in triplicate, and the corresponding pH buffer containing 10% (v/v) DMSO was used as the respective blank. Before measurement, the plate was equilibrated at 25 °C for 30 min. The absorption spectral measurements were performed in a plate reader (CLARIOstar® Plus, BMG Labtech, Germany) operated with CLARIOstar® software (5.70 R2) and ten repetitive measurements were performed and averaged to reduce instrument noise. Before each repetition, the plate was shaken for 30 s at 300 rpm, and each well was scanned with four flashes. The obtained absorption spectra were further processed in MARS software (3.42 R5; BMG

Labtech), analyzed and plotted with OriginPro 2019 (9.6.0.172, OriginLab Corporation, USA).

**Confocal and STED (stimulated emission depletion) microscopy**

Confocal and STED images were acquired using two Abberior Expert Line (Abberior Instruments GmbH, Göttingen, Germany) fluorescence microscopes built on a motorized inverted microscope IX83 (Olympus, Tokyo, Japan). Microscope 1 is equipped with pulsed STED lasers at 595 nm and 775 nm shaped by Spatial Light Modulators (SLMs), and with 355 nm, 405 nm, 485 nm, 561 nm, and 640 nm excitation lasers, and a 100x/1.40 oil immersion objective lenses (Olympus). Microscope 2 is equipped with pulsed STED lasers at 655 nm and 775 nm, and with 520 nm, 561 nm, 640 nm, and multiphoton (Chameleon Vision II, Coherent, Santa Clara, USA) excitation lasers, and a 60x/1.42 oil immersion objective lens (Olympus). The multiphoton laser is tuneable in the 680 nm – 1080 nm range. Spectral detection is performed in both cases with avalanche photodiodes at spectral windows adjusted for each particular fluorophore.

Imaging and image processing was done with ImSpector software (v. 16.3.13367; Abberior Instruments GmbH, Göttingen, Germany), and all images are displayed as raw data unless otherwise noted.

Particular imaging conditions are given in Supplementary Table 1.

**Superresolution single molecule localization microscopy (SMLM) / Photoactivated localization microscopy (PALM)**

Images were acquired on a custom-built setup [2], equipped with a 473 nm (500 mW), a 532 nm (1 W) and a 560 nm (1 W) laser for excitation, a 405 nm (300 mW) laser for activation, a back illuminated EMCCD camera (Andor iXon 897 / 512×512 sensor, operated with an EM gain 200), and a Leica HCX PL APO CS 100x/1.46 oil lens. Emission light was separated from the excitation and activation light with proper combination of a dichroic mirror (Brightline HC 560, Semrock Inc., Rochester, NY, USA) and an emission filter (ET585/40 M, Chroma Technology Corp, Bellow Falls, VT,

USA). A movable mirror was used to switch between wide field, highly inclined and laminated optical sheet (HILO) and total internal reflection fluorescence (TIRF) illumination modes. Images were acquired with a 20 ms exposure time, and actual excitation laser powers in the back focal plane of approx. 250 mW for 473 nm. The 405 nm activation laser was incorporated as 300 μs pulses, in between frames, with a power of 0.001-1 mW in the back focal plane. The microscope components were controlled with custom LabView (2019 32bit) software, and the camera was controlled with the Andor Solis 4.31.30022 software package.

All images were analyzed and processed using the ThunderSTORM plugin [4] on ImageJ (version 1.52p). In brief, images were filtered with a wavelet filer (B-spline order 3, scale 2.0), approximate localization of the molecules was performed with a local maximum method (peak intensity threshold of ca. 2-3 standard deviations; connectivity 8-neighbourhood), and sub-pixel localization of the molecules was performed with a maximum likelihood fitting method (PSF integrated method, fitting radius 3 px, initial sigma 1.3–1.6). Post-processing was performed with the same plug-in. Data was drift-corrected based on the cross-correlation method, merged, and a density filter was applied (distance radius approx. 50 nm, minimum neighbors in the radius ca. 2-5). Filtering and rendering of localizations was done using custom-built MatLab (version R2007a) routines [2]. Final images were produced using an amplitude-normalized Gaussian rendering method with a fixed sigma value corresponding to the average localization uncertainty (25 nm) and a pixel-size of 2 nm for the rendered image.

**Labeling of antibodies**

Amino-reactive NHS-esters of dyes were coupled to secondary antibodies (Jackson ImmunoResearch Europe Ltd. product nos.: 111-005-003 or 115-005-003) using a standard coupling protocol. In brief, the reactive dye (**1-NHS**, **2-NHS, 3-NHS** or DyLight 515-LS NHS ester (ThermoFisher 82491) was dissolved in anhydrous DMSO (2 mg $ml^{-1}$), and mixed with 0.5 mg antibody in a proportion of 5–10 equivalents (dye/protein). The pH of the solution was adjusted to ≈8.4 with carbonate buffer (1 M), and stirred for 1 h at RT while protected from ambient light. The mixture was purified using PD MiniTrap G-25 column (GE Healthcare). UV-Vis measurements in a small volume spectrometer (DS-11+, DeNovix) were performed to determine the fractions containing the protein, and for the determination of the degree of labelling (DOL) which ranged from 3.8 to 8.2.

#### Cell culture

COS-7, U2OS-Vim-Halo [5], U2OS-Vim-SNAP [5], and U2OS-NUP96-Halo [6] were cultured in Dulbecco's Modified Eagle Medium (DMEM, 4.5 g/L glucose) containing GlutaMAX and sodium pyruvate (ThermoFisher 31966), supplemented with 10 % (v/v) fetal bovine serum (FBS, ThermoFisher 10500064) and 1% Pen Strep (GIBCO 15140122) in a humidified 5 % CO2 incubator at 37 °C. Cells were split every 2 – 4 days or at confluency and regularly tested for mycoplasma contamination.

#### Indirect immunofluorescence labeling with antibody conjugates

Cells were grown for 24–48 h on glass coverslips and washed twice with PBS (pH 7.4), and fixed with either methanol (MeOH), preheated (37°C) paraformaldehyde (PFA), or glutaraldehyde (GLA) – depending on the favored method for the chosen antibodies or the imaging structures.

For MeOH fixation (Figure S1), the samples were treated with MeOH previously cooled to -20°C for 4 min and washed twice with PBS.

For PFA fixation for preferential preservation of mitochondria [7] (Figure 3), the samples were treated with an 8% formaldehyde solution in PBS 37 °C for 5 min, washed twice with PBS, and permeabilized in 0.5% Triton X-100 in PBS for 15 min at RT.

For GLA fixation of microtubules (Figure 2b,c and Figures S4–S6) and samples for four color imaging (Figure 4 and Figure S9a), samples were pre-permeabilized with 0.3% (v/v) glutaraldehyde (GLA) supplemented with 0.25% (v/v) Triton X-100 in cytoskeleton buffer (CB, 10 mM MES, 150 mM NaCl, 5 mM EGTA, 5 mM Glucose, 5 mM $MgCl_2·6H_2O$ pH 6.1) [8] for 2 min and fixed in 2% GLA in CB for 10 min. Samples were quenched in 0.1% (w/v) sodium borohydride for 7 min and washed three times with PBS.

To reduce unspecific binding, blocking buffer (2% or 5% BSA in PBS) was added and incubated for 1 h at room temperature. For samples fixed with PFA, Triton X-100 was added to the blocking buffer to a concentration of 0.5% and blocking was performed for 15 min at RT.

The coverslips were overlaid with the primary antibody solution in blocking buffer and incubated in a humid chamber for 1 h at room temperature, or overnight at 4 °C, and then washed with blocking buffer (3×5 min). The coverslips were then incubated with the secondary antibody in blocking buffer, in a humid chamber for 1 h at room temperature, and then washed with blocking buffer (3×5 min), and with PBS (2×5 min). After labelling with photoactivatable conjugates, the samples were protected from the ambient light. For four-color imaging, samples were further incubated with SiR-Actin [9] (500 nM in PBS, overnight at 4 °C; cat. # SC001, Spirochrome SA, Stein am Rhein, Switzerland) and DAPI (200 µL of 100 ng/mL) for 5 min at RT.

Samples were mounted with PBS or Mowiol supplemented with DABCO and sealed with nail polish.

Details on antibodies used can be found in the Supplementary Table S2.

**Optical microscopy of cells with lysosome-specific label 3-Lyso**

A stock solution of the Pepstatin A derivative of compound **3** (**3-Lyso**) was prepared in DMSO (approx. 1 mM). U2OS cells were grown for 12–72 h on glass coverslips. The cells were washed twice with PBS and incubated overnight in labeling medium (DMEM FluoroBrite medium supplemented with GlutaMAX (GIBCO 35050-038), 10 % (v/v) fetal bovine serum (FBS, ThermoFisher 10500064) and 1% Pen Strep (GIBCO 15140122) containing 20 nM **3-Lyso**. The medium was replaced with fresh medium containing 500 nM SiR-lysosome Kit (Spirochrome) and incubated for additional 60 min, before replacing with fresh medium for imaging.

Colocalization analysis (Pearson r) was performed in Coloc 2 plugin on ImageJ (version 1.52p).

**Optical microscopy cells with self-labelling enzymes**

Stock solutions of HaloTag ligand (**3-Halo**) or SNAP-tag ligand (**3-BG**) derivatives of large Stokes shift dye **3** were prepared in DMSO (1 mM). U2OS cells that stably expressed Vimentin-HaloTag, Vimentin-SNAP [5], or NUP96-HaloTag [6] were grown for 24–48 h on glass coverslips. Cells were incubated in the dark for 1 h (**3-BG**) or overnight (**3-Halo**) with the respective fluorescent ligands diluted from DMSO stock solutions with labeling medium to a final concentration of 200 nM. After labeling with ligands, the samples were protected from the ambient light. Cells were washed with labeling medium for 1 h; then the medium was changed again for fresh medium for live-cell imaging. For four-color experiments, the cells were co-stained with the always-on dyes MitoTracker Deep Red (10 nM), Hoechst (1 µl or 10 mg $ml^{-1}$) (Hoechst 33342 Solution, ThermoFischer Scientific), LIVE 510-tubulin (2 µM, Abberior GmbH) before the final wash.

Fixation after live labelling, for preservation of nuclear pore complexes (Figure S7), was performed with a 4% formaldehyde solution in PBS at room temperature for 25 min, washed once with a quenching solution (QS, 0.1 M $NH_4Cl$ and 0.1 M glycine in PBS) and then incubated with QS for 10 min at room temperature. Samples were then washed with PBS (2×5 min) and mounted in PBS for PALM imaging.

**Optical microscopy of cells with 3-Tz**

Stock solutions of bicyclo[6.1.0]non-4-yne (BCN) HaloTag ligand (HTL-BCN, prepared according to [10]) and tetrazine-functionalized large Stokes shift derivative (**3-Tz**) were prepared in DMSO (10 and 1 mM, respectively). U2OS cells that stably expressed Vimentin-HaloTag were grown for 24–48 h on glass coverslips.

Cells were incubated for 30 min with HTL-BCN diluted from DMSO stock solution with labeling medium to a final concentration of 10 µM [11]. Cells were then incubated in the dark overnight with the **3-Tz** diluted from DMSO stock solution with labeling medium to a final concentration of 200 nM. After labeling with ligands, the samples were protected from the ambient light. Cells were washed with fresh labelling medium for 1 h; then the medium was changed for fresh medium for live-cell imaging.

##### Single detection channel two-color multiplexing by photoactivation

PFA-fixed COS-7 cells were prepared by indirect immunofluorescence labelling of microtubules with DyLight 515-LS and mitochondria labelled with compound **3-NHS** as described above. Sequential imaging was performed on a confocal setup before photoactivation, after photobleaching of DL515-LS with high power of 485 nm light, and after photoactivation of **3** with 405 nm activation. All images were acquired with the same excitation laser, 485 nm, and detection channel, 560 nm–700 nm range. A single-channel, pseudo two-color (multiplexed) image of two different targets may be obtained from overlaying the images.

#### Synthesis of photoactivatable fluorescent dyes and model compounds

##### Synthesis of dye 1

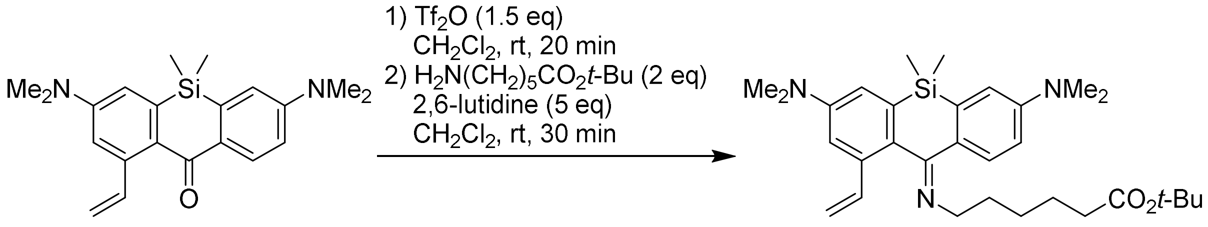

**S-1** **S-2**

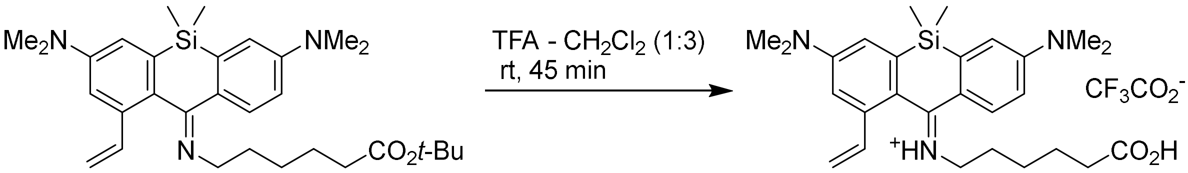

**S-2** **1**

Compound **S-1** (20 mg, 0.057 mmol; compound 1 in [2]) was dissolved in dichloromethane (500 µL). Trifluoromethanesulfonic anhydride solution (86 µl of 1 M in dichloromethane, 0.086 mmol, 1.5 equiv) was added dropwise, and the solution was stirred for 20 min at rt. The resulting blue solution was transferred dropwise to a stirring solution of *tert*-butyl 6-aminohexanoate (21.3 mg, 0.114 mmol, 2 equiv) and 2,6-lutidine (30.5 mg, 0.285 mmol) in dichloromethane (500 µL), cooled in an ice-water bath. An additional rinse of dichloromethane (500 µL) was used to ensure complete transfer. The solution was stirred for 30 min, then sat. aq. $NaHCO_3$ (10 mL) was added and the reaction mixture was extracted with ethyl acetate (3 × 10 mL). The combined extracts were washed with brine (50 mL), dried over $Na_2SO_4$, filtered, evaporated, the product was isolated by flash chromatography on a Biotage Isolera system (12g Interchim SiHP 30 µm cartridge, 20 to 100% ethyl acetate/hexane) and freeze-dried from 1,4-dioxane to yield 20 mg (66%) of the intermediate **S-2** as a brown oil which was used in the following step without further characterization.

To a solution of compound **S-2** (20 mg, 0.038 mmol) in dichloromethane (300 µL), trifluoroacetic acid (100 µL) was added dropwise. The reaction mixture was stirred for 45 minutes at rt, protected from light. The volatiles were removed *in vacuo* by coevaporation with toluene (3 × 10 ml) and freeze-dried from dioxane to yield 22 mg (~100%, or 66% over 2 steps) of **1** as an orange solid (mixture of *(E)*- and *(Z)*-isomers of the imine in 2:1 ratio).

$^1H$ NMR, major *(E)*-isomer (400 MHz, DMSO-$d_6$): δ 11.99 (br.s, 1H), 11.54 (d, *J* = 8.1 Hz, 1H), 7.76 (d, *J* = 8.8 Hz, 1H), 7.06 (d, *J* = 2.6 Hz, 1H), 7.04 (d, *J* = 2.7 Hz, 1H), 6.96 (d, *J* = 2.6 Hz,

1H), 6.91 (dd, $J$ = 8.8, 2.7 Hz, 1H), 6.74 (dd, $J$ = 17.3, 10.9 Hz, 1H), 5.88 (dd, $J$ = 17.3, 0.9 Hz, 1H), 5.45 (dd, $J$ = 10.9, 0.9 Hz, 1H), 3.40 – 3.20 (m, 2H), 3.10 (s, 6H), 3.07 (s, 6H), 2.05 (t, $J$ = 7.3 Hz, 2H), 1.65 – 1.52 (m, 2H), 1.36 – 1.18 (m, 4H), 0.68 (s, 3H), 0.32 (s, 3H).

$^{13}$C NMR, major *(E)*-isomer (101 MHz, DMSO-$d_6$): δ 175.5, 173.8, 151.3, 150.9, 139.4, 137.7, 137.6, 134.5, 128.6, 125.6, 124.6, 118.3, 115.8, 115.3, 111.9, 109.0, 49.0, 39.4, 39.3, 32.9, 26.9, 25.2, 23.4, -0.5, -5.2.

HRMS ($C_{27}H_{37}N_3O_2Si$): *m/z* (positive mode) = 464.2725 (found $[M+H]^+$), 464.2728 (calc.).

#### Synthesis of dye 2 and related model compound 2a

##### Compound 2a

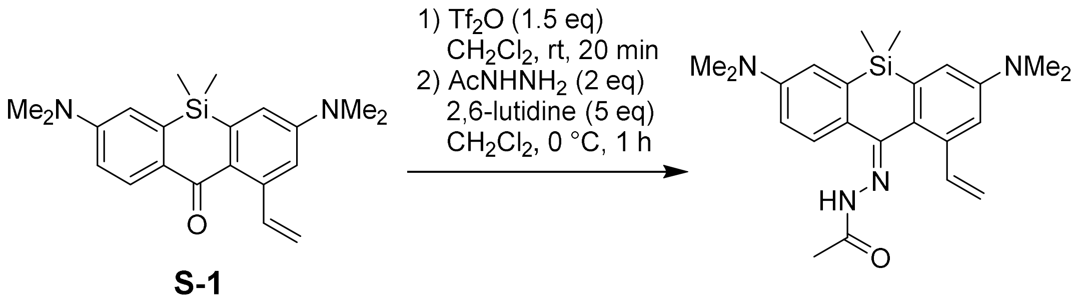

A solution of trifluoromethanesulfonic anhydride ($Tf_2O$, 1 M in $CH_2Cl_2$; 0.15 mL, ~0.15 mmol, 1.5 equiv.) was added to the stirred solution of **S-1** [2] (35 mg, 0.1 mmol) in dry $CH_2Cl_2$ (2 mL) under argon, and the resulting dark blue solution was stirred at rt for 20 min. It was then transferred dropwise into the stirred mixture of acethydrazide (15 mg, 0.2 mmol, 2 equiv.), 2,6-lutidine (58 µL, 0.5 mmol, 5 equiv.) and dry $CH_2Cl_2$ (1 mL), cooled in ice-water bath. The reaction mixture was stirred at 0 °C for 1 h, poured into sat. aq. $NaHCO_3$ (50 mL), the product was then extracted with ethyl acetate (3×25 mL) and the combined extracts were dried over $Na_2SO_4$. The product was isolated by flash column chromatography (12 g Interchim SiHP 30 µm cartridge, gradient 0% to 50% EtOAc/$CH_2Cl_2$) and freeze-dried from 1,4-dioxane to yield 23 mg (57%) of **2a** as a mixture of (*E/Z*)-stereoisomers (ratio 0.55:0.45; purity 90%).

$^1$H NMR (400 MHz, $CDCl_3$): δ 8.98 (s, 0.55H, major), 8.40 (s, 0.45H, minor), 7.63 (d, $J$ = 8.6 Hz, 0.45H, minor), 7.42 (d, $J$ = 8.6 Hz, 0.55H, major), 7.27 (dd, $J$ = 17.5, 10.9 Hz, 0.55H, major), 6.98 – 6.91 (m, 2×0.55+0.45H, major+minor), 6.88 – 6.83 (m, 2×0.45+0.55H, major+minor), 6.80 (dd, $J$ = 8.6, 2.7 Hz, 0.45H, minor), 6.72 (dd, $J$ = 8.6, 2.8 Hz, 0.55H, major), 6.54 (dd, $J$ = 17.5, 10.9 Hz, 0.45H, minor), 5.80 (dd, $J$ = 17.5, 1.0 Hz, 0.45H, minor), 5.64 (dd, $J$ = 17.5, 1.5 Hz, 0.55H, major), 5.30 (dd, $J$ = 11.0, 1.0 Hz, 0.45H, minor), 5.23 (dd, J = 10.9, 1.5 Hz, 0.55H, major), 3.03 (s, 6×0.45H, minor), 3.01 (s, 6×0.55H, major), 3.01 (s, 6×0.55H, major), 2.99 (s, 6×0.45H, minor), 2.34 (s, 3×0.55H, major), 2.34 (s, 3×0.45H, minor), 0.61 (s, 3×0.45H, minor), 0.28 (s, 3×0.55H, major).

$^{13}$C NMR (101 MHz, $CDCl_3$): δ 173.6, 173.0, 150.24, 150.15, 149.97, 149.95, 149.7, 148.5, 139.7, 139.6, 138.2, 137.8, 136.65, 136.61, 135.7, 135.2, 134.3, 131.5, 127.4, 126.6, 125.1, 124.4, 117.0, 116.8, 116.5, 116.1, 115.6, 113.7, 113.5, 112.5, 111.7, 109.7, 40.74, 40.70, 40.5, 40.4, 20.8, 20.6, -0.2, -4.9.

HRMS ($C_{23}H_{30}N_4OSi$): *m/z* (positive mode) = 407.2256 (found $[M+H]^+$), 407.2262 (calc.).

**Compound 2a-CF**

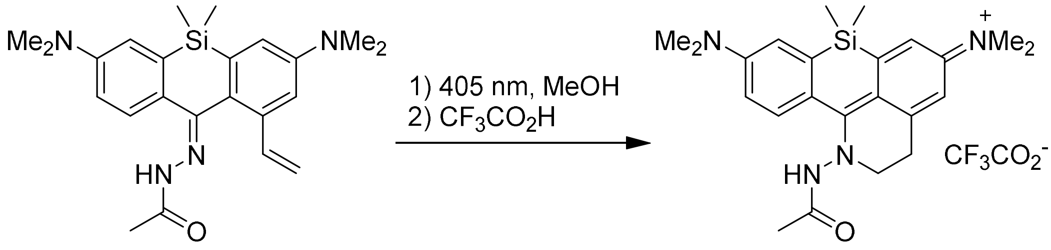

**2a** **2a-CF**

Compound **2a** (10 mg, 24.6 µmol) was placed in a 10 mL pear-shaped flask equipped with a stirring bar, dissolved in degassed methanol (5 mL) and irradiated in a Penn OC Photoreactor m1 using a custom 405 nm LED light source (20% power) until the reaction was found complete by HPLC analysis (50 min total time). Trifluoroacetic acid (10 µL) was added, the resulting bright red solution was evaporated, the product was isolated by preparative HPLC (Interchim Uptisphere Strategy PhC4 250×21.2 mm 5 µm, solvent flow rate 18 mL/min, gradient 30% to 70% A:B, A – acetonitrile + 0.1% (v/v) trifluoroacetic acid, B – water + 0.1% (v/v) trifluoroacetic acid) and freeze-dried from aq. dioxane to give 10 mg (78%) of **2a-CF** as red solid.

$^{1}$H NMR (400 MHz, $CDCl_3$): 12.28 (br.s, 1H), 8.03 (d, *J* = 8.9 Hz, 1H), 6.83 (d, *J* = 2.6 Hz, 1H), 6.81 (d, *J* = 2.6 Hz, 1H), 6.76 – 6.68 (m, 1H), 6.49 (d, J = 2.6 Hz, 1H), 4.02 (br.s, 2H), 3.17 (s, 6H), 3.15 (br.s, 2H), 3.13 (s, 6H), 1.95 (s, 3H), 0.53 (s, 6H).

$^{13}$C NMR (101 MHz, $CDCl_3$): δ 169.8, 166.7, 152.5, 151.9, 142.5, 142.3, 139.8, 133.1, 120.3, 119.9, 115.6, 115.4, 112.3, 110.6, 52.5, 40.2, 40.0, 28.7, 20.6.

$^{19}$F NMR (376 MHz, $CDCl_3$): δ -75.7 ($CF_3CO_2^-$).

HRMS ($C_{23}H_{31}N_4OSi$): *m/z* (positive mode) = 407.2258 (found $[M]^+$), 407.2262 (calc.).

#### Compound S-4

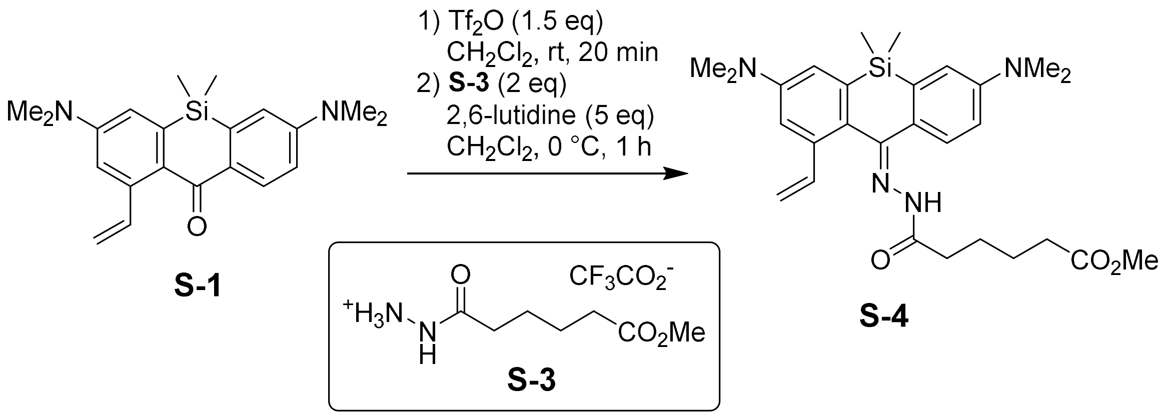

A solution of trifluoromethanesulfonic anhydride ($Tf_2O$, 1 M in $CH_2Cl_2$; 0.15 mL, ~0.15 mmol, 1.5 equiv.) was added to the stirred solution of **S-1** (35 mg, 0.1 mmol) in dry $CH_2Cl_2$ (2 mL) under argon, and the resulting dark blue solution was stirred at rt for 20 min. It was then transferred dropwise into the stirred mixture of **S-3** (compound "Me-Hydrazine" in [12]; 58 mg, 0.2 mmol, 2 equiv.), 2,6-lutidine (58 µL, 0.5 mmol, 5 equiv.) and dry $CH_2Cl_2$ (1 mL), cooled in ice-water bath. The reaction mixture was stirred at 0 °C for 1 h, poured into sat. aq. $NaHCO_3$ (50 mL), the product was then extracted with ethyl acetate (3×25 mL) and the combined extracts were dried over $Na_2SO_4$. The product was isolated by flash column chromatography (12 g Interchim SiHP 30 µm cartridge, gradient 0% to 50% EtOAc/$CH_2Cl_2$) and freeze-dried from 1,4-dioxane to yield 30 mg (59%) of **S-4** as a mixture of (*E/Z*)-stereoisomers (ratio 0.57:0.43; each existing in the form of a pair of *(E/Z)* amide bond rotamers – interpreted whenever possible in the $^1H$ NMR data below).

$^1H$ NMR (400 MHz, $CD_3CN$): δ 9.15 (s, 0.10H, major isomer/minor rotamer), 8.84 (s, 0.47H, major isomer/major rotamer), 8.61 (s, 0.15H, minor isomer/minor rotamer), 8.23 (s, 0.28H, minor isomer/major rotamer), 7.57 (d, *J* = 8.6 Hz, 0.28H, minor isomer/major rotamer), 7.54 (d, *J* = 8.6 Hz, 0.16H, minor isomer/minor rotamer), 7.45 (d, *J* = 8.6 Hz, 0.57H, major isomer/major+minor rotamers), 7.41 (dd, *J* = 17.8, 10.7 Hz, 0.10H, major isomer/minor rotamer), 7.27 (dd, *J* = 17.4, 10.9 Hz, 0.47H, major isomer/major rotamer), 7.05 (dd, *J* = 7.5, 2.7 Hz, 1H, major+minor isomer/major+minor rotamer), 6.99 – 6.90 (m, major+minor isomer/major+minor rotamer, 2H), 6.85 – 6.75 (m, 1H, major+minor isomer/major+minor rotamer), 6.58 – 6.45 (m, 0.43H, minor isomer/major+minor rotamer), 5.93 – 5.83 (m, 0.43H, minor isomer/major+minor rotamers), 5.76 – 5.63 (m, 0.57H, major isomer/major+minor rotamers), 5.31 – 5.23 (m, 0.43H, minor isomer/major+minor rotamers), 5.24 – 5.17 (m, 0.57H, major isomer/major+minor rotamers), 3.60 (s, 3H), 3.05 – 3.01 (m, 3H), 3.00 – 2.98 (m, 6H), 2.96 (s, 3H), 2.85 – 2.49 (m, 2H), 2.38 – 2.24 (m, 2H), 1.70 – 1.51 (m, 4H), 0.63 (s, 1.29H, minor isomer/major+minor rotamer), 0.62 (s, 1.71H, major isomer/major+minor rotamer), 0.21

(s, 1.29H, minor isomer/major+minor rotamer), 0.20 (s, 1.71H, major isomer/major+minor rotamer).

$^{13}$C NMR (101 MHz, $CD_3CN$): δ 175.3, 174.7, 174.6, 174.5, 169.2, 153.2, 151.24, 151.17, 150.9, 150.8, 150.6, 150.3, 148.6, 140.5, 140.4, 140.2, 139.1, 138.7, 137.2, 137.0, 136.6, 135.8, 135.5, 134.9, 132.3, 128.1, 127.6, 127.4, 125.9, 125.6, 124.9, 118.1, 117.7, 117.0, 116.5, 116.3, 116.2, 114.2, 114.1, 113.5, 113.2, 111.6, 110.0, 109.8, 51.90, 51.88, 40.7, 40.64, 40.61, 40.43, 40.36, 35.3, 34.27, 34.26, 34.2, 33.0, 32.9, 30.6, 25.6, 25.4, 25.3, 25.2, 25.02, 25.00, -0.4, -5.1.

HRMS ($C_{28}H_{38}N_4O_3Si$): *m/z* (positive mode) = 507.2776 (found $[M+H]^+$), 507.2786 (calc.).

**Dye 2**

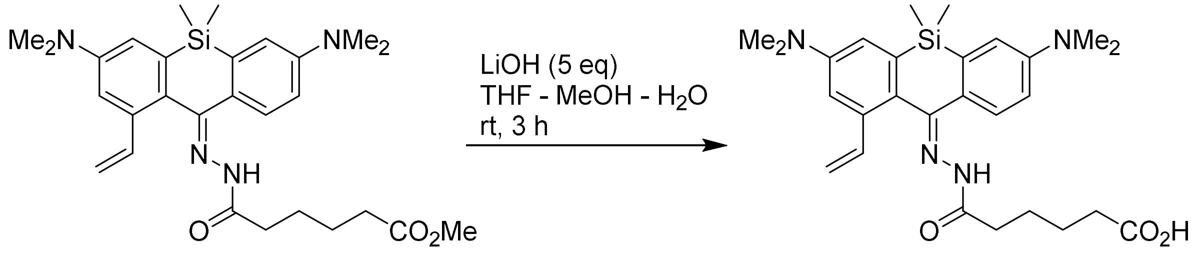

**S-4** **2**

A solution of lithium hydroxide monohydrate (12.5 mg, 0.30 mmol, 5 equiv.) in water (0.5 mL) was added to the solution of **S-4** (30 mg, 59.2 µmol) in the mixture of THF (2 mL) and methanol (0.5 mL), and the reaction mixture was stirred at rt for 3 h. The organic solvents were removed on a rotary evaporator, and the product was isolated by preparative HPLC (Interchim Uptisphere Strategy PhC4 250×21.2 mm 5 µm, solvent flow rate 18 mL/min, gradient 20% to 70% A:B, A – acetonitrile + 0.1% (v/v) $HCO_2H$, B – water + 0.1% (v/v) $HCO_2H$) and freeze-dried from aq. dioxane to give 25 mg (86%) of **2** as orange-brown solid (mixture of (*E/Z*)-stereoisomers in ratio 0.66:0.34, with the minor stereoisomer existing in the form of a pair of *(E/Z)* amide bond rotamers).

$^1$H NMR (400 MHz, $CDCl_3$): δ 9.02 (s, 0.20H, minor isomer/major rotamer), 8.42 (s, 0.80H, major isomer + minor isomer/minor rotamer), 7.80 (d, *J* = 8.5 Hz, 0.14H, minor isomer/minor rotamer), 7.62 (d, *J* = 8.5 Hz, 0.66H, major isomer), 7.41 (d, *J* = 8.6 Hz, 0.20H, minor isomer/major rotamer), 7.24 (dd, *J* = 17.5, 10.9 Hz, 0.20H, minor isomer/major rotamer), 6.99 – 6.73 (m, 4H), 6.58 (dd, *J* = 17.4, 10.9 Hz, 0.14H, minor isomer/minor rotamer), 6.52 (dd, J = 17.4, 10.9 Hz, 0.66H, major isomer), 5.84 (d, *J* = 18.1 Hz, 0.14H, minor isomer/minor rotamer), 5.79 (dd, *J* = 17.5, 1.0 Hz, 0.66H, major isomer), 5.65 (dd, *J* = 17.4, 1.4 Hz, 0.20H, minor isomer/major rotamer), 5.29 (dd, *J* = 10.9, 1.0 Hz, 0.80H, major isomer + minor isomer/minor

rotamer), 5.24 (dd, $J$ = 10.9, 1.4 Hz, 0.20H, minor isomer/major rotamer), 3.07 (s, 0.7H), 3.03 (s, 4.3H), 3.02 (s, 2H), 3.00 (s, 4H), 2.98 (s, 1H), 2.93 – 2.62 (m, 2H), 2.46 – 2.30 (m, 2H), 1.85 – 1.62 (m, 4H), 0.61 (s, 3H), 0.28 (s, 3H).
$^{13}C$ NMR (101 MHz, $CDCl_3$): δ 178.4, 178.2, 175.7, 175.1, 168.4, 150.2, 149.9, 148.7, 139.64, 139.56, 138.1, 136.7, 136.3, 135.6, 135.3, 127.5, 126.6, 124.2, 117.0, 116.8, 116.6, 115.8, 113.8, 112.6, 109.7, 40.8, 40.5, 40.4, 35.4, 33.8, 33.7, 32.5, 32.4, 24.9, 24.6, 24.52, 24.48, 24.3, 24.2, -0.2, -5.0.
HRMS ($C_{27}H_{36}N_4O_3Si$): $m/z$ (positive mode) = 493.2626 (found $[M+H]^+$), 493.2629 (calc.).

##### Synthesis of dye 3 and related model compound 3a

##### Compound S-5

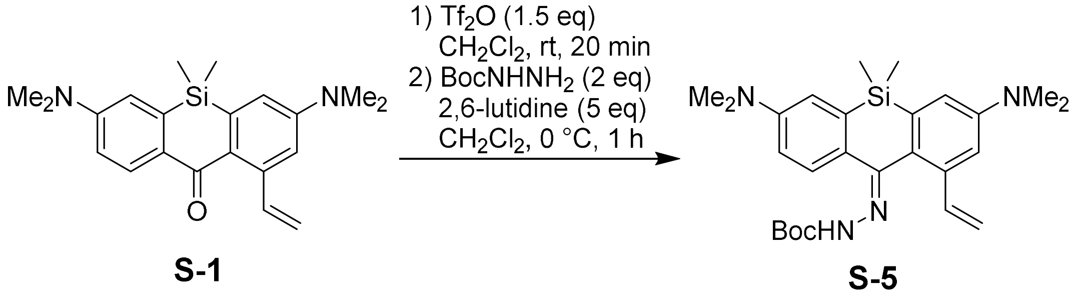

A solution of trifluoromethanesulfonic anhydride ($Tf_2O$, 1 M in $CH_2Cl_2$; 0.34 mL, ~0.34 mmol, 1.5 equiv.) was added to the stirred solution of **S-1** [2] (80 mg, 0.23 mmol) in dry $CH_2Cl_2$ (3 mL) under argon, and the resulting dark blue solution was stirred at rt for 20 min. It was then transferred dropwise into the stirred mixture of *tert*-butyl carbazate (61 mg, 0.46 mmol, 2 equiv.), 2,6-lutidine (133 µL, 1.14 mmol, 5 equiv.) and dry $CH_2Cl_2$ (2 mL), cooled in ice-water bath. The reaction mixture was stirred at 0 °C for 1 h, poured into sat. aq. $NaHCO_3$ (50 mL), the product was then extracted with ethyl acetate (3×25 mL) and the combined extracts were dried over $Na_2SO_4$. The product was isolated by flash column chromatography (25 g Interchim SiHP 30 µm cartridge, gradient 5% to 30% EtOAc/hexane + 20% $CH_2Cl_2$ constant additive) and freeze-dried from 1,4-dioxane to yield 63 mg (59%) of **S-5** as a mixture of (*E/Z*)-stereoisomers (ratio 0.70:0.30).

$^1H$ NMR (400 MHz, $CD_3CN$): δ 8.31 (s, 0.7H, major), 7.76 (s, 0.3H, minor), 7.50 (d, $J$ = 8.6 Hz, 0.3H, minor), 7.42 (d, $J$ = 8.6 Hz, 0.7H, major), 7.36 (dd, $J$ = 17.6, 11.0 Hz, 0.7H, major), 7.06 (d, $J$ = 2.7 Hz, 0.7H, major), 7.04 (d, $J$ = 2.6 Hz, 0.3H, minor), 6.97 – 6.95 (m, 1H, major+minor), 6.94 – 6.91 (m, 1H, major+minor), 6.82 – 6.77 (m, 1H, major+minor), 6.55 (dd, $J$ = 17.5, 10.9 Hz, 0.3H, minor), 5.88 (dd, $J$ = 17.5, 1.1 Hz, 0.3H, minor), 5.69 (dd, $J$ = 17.6, 1.4 Hz, 0.7H, major), 5.31 (dd, $J$ = 10.9, 1.1 Hz, 0.3H, minor), 5.19 (dd, $J$ = 11.0, 1.4 Hz, 0.7H, major), 3.03

(s, 6×0.3H, minor), 2.99 (s, 6×0.7H, major), 2.98 (s, 6×0.7H, major), 2.96 (s, 6×0.3H, minor), 1.45 (s, 9×0.7H, major), 1.42 (s, 9×0.3H, minor), 0.62 (s+br.s, 3H, major+minor), 0.21 (s+br.s, 3H, major+minor).

$^{13}$C NMR (101 MHz, $CD_3CN$): δ 153.6, 153.4, 151.3, 151.1, 150.9, 150.8, 150.5, 149.6, 140.5, 140.3, 138.7, 138.2, 137.2, 136.7, 135.5, 135.0, 132.6, 127.8, 127.4, 126.5, 126.4, 125.4, 117.8, 117.1, 116.6, 116.3, 114.2, 113.9, 113.2, 111.1, 109.8, 80.93, 80.86, 40.68, 40.66, 40.5, 40.4, 28.5, -0.4, -5.1.

HRMS ($C_{23}H_{30}N_4OSi$): *m/z* (positive mode) = 407.2256 (found $[M+H]^+$), 407.2262 (calc.).

**Compound 3a**

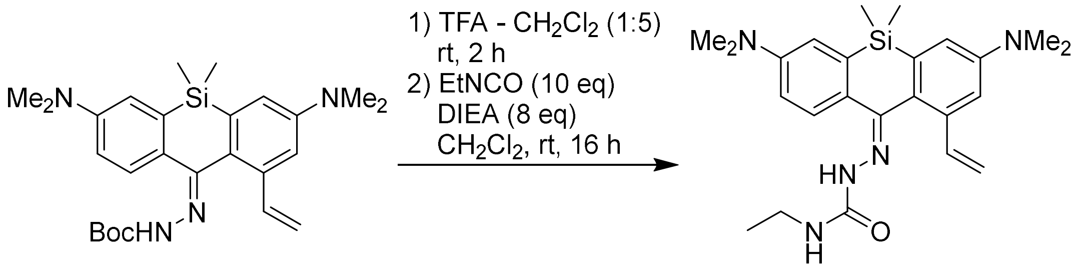

**S-5** **3a**

Compound **S-5** (35 mg, 75.3 μmol) was dissolved in the mixture of trifluoroacetic acid (0.2 mL) and $CH_2Cl_2$ (1 mL) and stirred at rt for 2 h. Toluene (2 mL) was added, the solvents were evaporated and the residue was chased with toluene-$CH_2Cl_2$ (2×2 mL). The residue was then dissolved in dry $CH_2Cl_2$ (1 mL), ethyldiisopropylamine (DIEA; 100 μL, 0.6 mmol, 8 equiv.) followed by ethyl isocyanate (59 μL, 0.75 mmol, 10 equiv.) were added, and the resulting solution was stirred at rt overnight (16 h). The reaction mixture was evaporated on Celite, the product was isolated by flash column chromatography (12 g Interchim SiHP 30 μm cartridge, gradient 0% to 50% EtOAc/$CH_2Cl_2$) and freeze-dried from 1,4-dioxane to yield 22 mg (67%) of **3a** as light yellow solid as a mixture of (*E/Z*)-stereoisomers (ratio 0.70:0.30).

$^{1}$H NMR (400 MHz, $CD_3CN$): δ 7.99 (s, 0.3H, minor), 7.58 (d, *J* = 8.6 Hz, 0.7H, major), 7.55 (d, *J* = 8.8 Hz, 0.3H, minor), 7.40 (s, 0.7H, major), 7.26 (dd, *J* = 17.5, 10.9 Hz, 0.3H, minor), 7.06 (d, *J* = 2.8 Hz, 0.3H, minor), 7.04 (d, *J* = 2.7 Hz, 0.7H, major), 6.98 (d, *J* = 2.7 Hz, 0.7H, major), 6.94 – 6.90 (m, 0.7+2×0.3H, major+minor), 6.85 – 6.78 (m, 1H, major+minor), 6.55 (dd, *J* = 17.5, 11.0 Hz, 0.7H, major), 6.39 (br.t, *J* = 6.0 Hz, 0.7H, major), 6.28 (br.t, *J* = 5.8 Hz, 0.3H, minor), 5.89 (dd, *J* = 17.4, 1.0 Hz, 0.7H, major), 5.65 (dd, *J* = 17.4, 1.6 Hz, 0.3H, minor), 5.28 (dd, *J* = 10.9, 1.0 Hz, 0.7H, major), 5.22 (dd, *J* = 10.9, 1.6 Hz, 0.3H, minor), 3.26 – 3.17 (m, 2H, major+minor), 3.02 (s, 6×0.7H, major), 3.00 (s, 6×0.3H, minor), 2.99 (s, 6×0.3H, minor),

2.96 (s, 6×0.7H, major), 1.11 (t, $J$ = 7.2 Hz, 3×0.3H, minor), 1.10 (t, $J$ = 7.2 Hz, 3×0.7H, major), 0.62 (s, 3H, major+minor), 0.20 (s, 3H, major+minor).

$^{13}C$ NMR (101 MHz, $CD_3CN$): δ 156.9, 156.4, 151.2, 150.90, 150.88, 150.6, 147.8, 145.7, 140.3, 140.1, 140.0, 138.4, 137.7, 137.1, 136.6, 135.4, 135.1, 132.3, 128.2, 127.6, 126.0, 125.5, 117.7, 116.9, 116.4, 116.3, 114.3, 113.5, 113.3, 112.4, 109.8, 40.74, 40.67, 40.5, 40.4, 35.1, 30.7, 21.6, 16.0, -0.4, -5.1.

HRMS ($C_{24}H_{33}N_5OSi$): $m/z$ (positive mode) = 436.2521 (found $[M+H]^+$), 436.2527 (calc.).

**Compound 3a-CF**

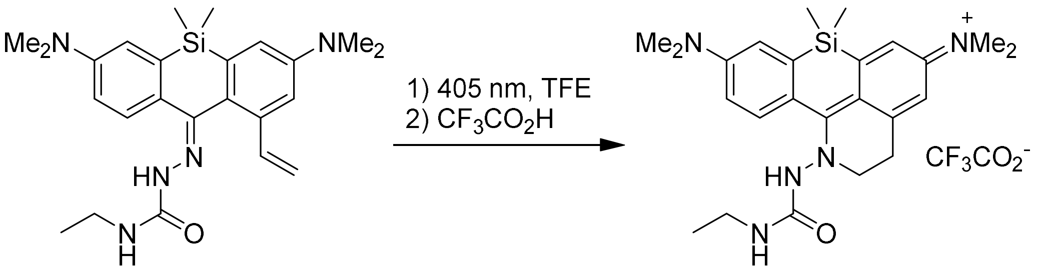

**3a** **3a-CF**

Compound **3a** (8.3 mg, 19.1 µmol) was placed in a 10 mL pear-shaped flask equipped with a stirring bar, dissolved in degassed TFE (2,2,2-trifluoroethanol; 5 mL) and irradiated in a Penn OC Photoreactor m1 using a custom 405 nm LED light source (20% power) until the reaction was found complete by HPLC analysis (20 min total time). The resulting bright red solution was evaporated, the product was isolated by preparative HPLC (Hypersil Gold C18 250×21.2 mm 5 µm, solvent flow rate 18 mL/min, gradient 20% to 70% A:B, A – acetonitrile + 0.1% (v/v) trifluoroacetic acid, B – water + 0.1% (v/v) trifluoroacetic acid) and freeze-dried from aq. dioxane to give 5.2 mg (50%) of **3a-CF** as red-orange fluffy solid.

$^1H$ NMR (400 MHz, $CDCl_3$): δ 11.01 (s, 1H), 8.09 (d, J = 9.2 Hz, 1H), 6.81 (app.t, J = 2.7 Hz, 2H), 6.70 (dd, J = 9.2, 2.7 Hz, 1H), 6.47 (d, J = 2.7 Hz, 1H), 4.45 (s, 3H, NH + $H_2O$), 4.05 (t, J = 7.0 Hz, 2H), 3.15 (s, 8H, 2× Me + $CH_2$), 3.12 (s, 8H, 2× Me + $CH_2$), 1.05 (t, J = 7.2 Hz, 3H), 0.52 (s, 3H), -0.00 (s, 3H).

$^{13}C$ NMR (101 MHz, $CDCl_3$): δ 169.9, 155.2, 152.3, 151.7, 142.4, 142.1, 139.5, 133.8, 120.8, 120.4, 115.4, 115.2, 112.2, 110.5, 53.4, 40.2, 40.0, 35.3, 28.9, 15.1, 0.1, -2.1.

$^{19}F$ NMR (376 MHz, $CDCl_3$): δ -75.7 ($CF_3CO_2^-$).

HRMS ($C_{30}H_{43}N_5O_3Si$): $m/z$ (positive mode) = 550.3203 (found $[M+H]^+$), 550.3208 (calc.).

#### Compound S-7

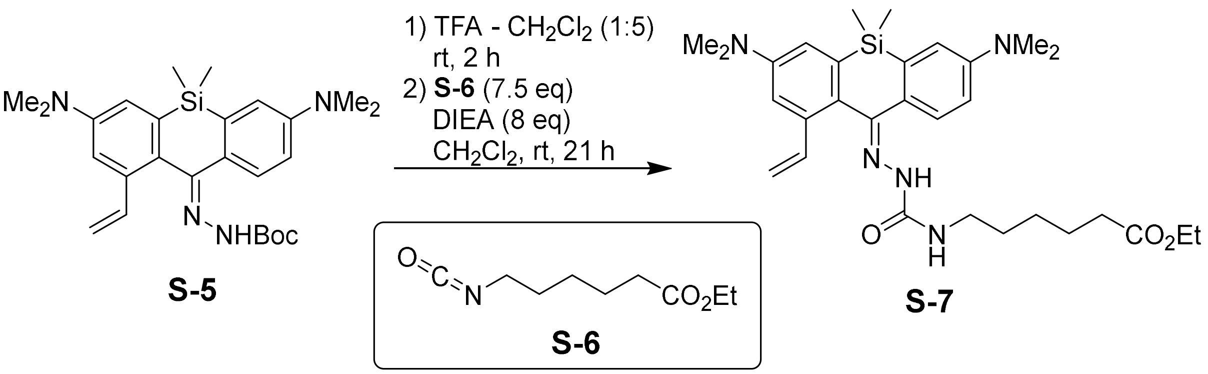

Compound **S5** (31 mg, 66.7 µmol) was dissolved in the mixture of trifluoroacetic acid (0.2 mL) and $CH_2Cl_2$ (1 mL) and stirred at rt for 2 h. Toluene (2 mL) was added, the solvents were evaporated and the residue was chased with toluene-$CH_2Cl_2$ (2×2 mL). The residue was then dissolved in dry $CH_2Cl_2$ (1 mL), ethyldiisopropylamine (DIEA; 100 µL, 0.54 mmol, 8 equiv.) followed by ethyl 6-isocyanatohexanoate (**S-6**; 90 µL, 0.51 mmol, 7.5 equiv.) were added, and the resulting solution was stirred at rt overnight (21 h). The reaction mixture was evaporated on Celite, the product was isolated by flash column chromatography (12 g Interchim SiHP 30 µm cartridge, gradient 0% to 50% EtOAc/$CH_2Cl_2$) and freeze-dried from 1,4-dioxane to yield 22 mg (60%) of **S-7** as yellow-brown viscous oil as a mixture of (*E/Z*)-stereoisomers (ratio 0.70:0.30).

$^{1}$H NMR (400 MHz, $CD_3CN$): δ 8.00 (s, 0.30H, minor isomer), 7.58 (d, *J* = 8.6 Hz, 0.70H, major isomer), 7.55 (d, *J* = 9.1 Hz, 0.30H, minor isomer), 7.40 (s, 0.70H, major isomer), 7.25 (dd, *J* = 17.5, 10.9 Hz, 0.30H, minor isomer), 7.06 (d, *J* = 2.8 Hz, 0.30H, minor isomer), 7.04 (d, *J* = 2.6 Hz, 0.70H, major isomer), 6.98 (d, *J* = 2.7 Hz, 0.70H, major isomer), 6.94 – 6.90 (m, 1.3H, major isomer + 2×minor isomer), 6.82 (dd, *J* = 8.7, 2.7 Hz, 0.30H, minor isomer), 6.80 (dd, *J* = 8.6, 2.8 Hz, 0.70H, major isomer), 6.54 (dd, *J* = 17.5, 11.0 Hz, 0.70H, major isomer), 6.40 (t, *J* = 6.1 Hz, 0.70H, major isomer), 6.30 (t, *J* = 5.9 Hz, 0.30H, minor isomer), 5.89 (dd, *J* = 17.5, 1.1 Hz, 0.70H, major isomer), 5.65 (dd, *J* = 17.4, 1.6 Hz, 0.30H, minor isomer), 5.27 (dd, *J* = 10.9, 1.1 Hz, 0.70H, major isomer), 5.21 (dd, *J* = 10.9, 1.6 Hz, 0.30H, minor isomer), 4.10 – 4.01 (m, 3H, major isomer + minor isomer), 3.23 – 3.13 (m, 2H, major isomer + minor isomer), 3.02 (s, 3×0.70H, major isomer), 2.99 (s, 3×0.30H, minor isomer), 2.99 (s, 3×0.30H, minor isomer), 2.96 (s, 3×0.70H, major isomer), 2.30 – 2.22 (m, 3H, major isomer + minor isomer), 1.63 – 1.42 (m, 6H, major isomer + minor isomer), 1.37 – 1.25 (m, 2H, major isomer + minor isomer), 1.23 – 1.15 (m, 6H, major isomer + minor isomer), 0.62 (s, 3H, major isomer + minor isomer), 0.20 (s, 3H, major isomer + minor isomer).

$^{13}$C NMR (101 MHz, $CD_3CN$): δ 174.3, 157.3, 156.9, 156.5, 151.2, 150.90, 150.87, 150.6, 147.9, 145.7, 140.3, 140.1, 140.0, 138.4, 137.6, 137.2, 136.6, 135.4, 135.1, 132.3, 128.2,

127.6, 126.0, 125.5, 117.7, 116.9, 116.4, 116.3, 114.3, 113.4, 113.3, 112.4, 109.8, 60.8, 55.3, 46.0, 40.9, 40.8, 40.74, 40.67, 40.50, 40.46, 40.4, 40.1, 34.8, 34.72, 34.71, 34.7, 30.8, 30.74, 30.65, 30.64, 30.1, 27.2, 27.06, 27.05, 26.99, 26.8, 25.5, 25.43, 25.40, 21.6, 14.6, -0.4, -5.1. HRMS ($C_{28}H_{38}N_4O_3Si$): *m/z* (positive mode) = 507.2776 (found $[M+H]^+$), 507.2786 (calc.).

**Dye 3**

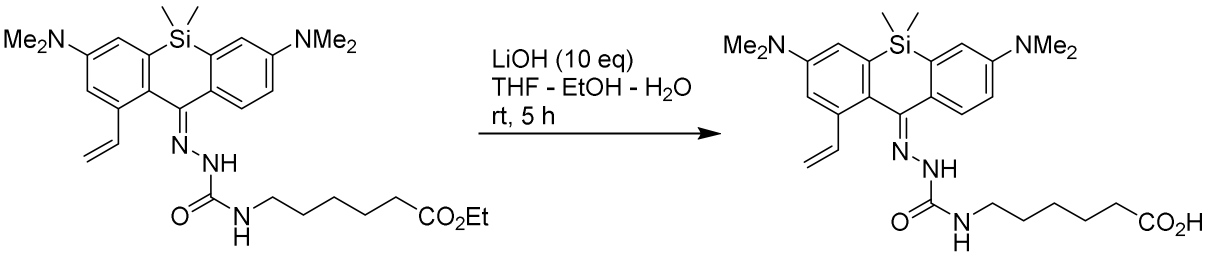

**S-7** **3**

A solution of lithium hydroxide monohydrate (16.8 mg, 0.40 mmol, 10 equiv.) in water (0.5 mL) was added to the solution of **S-7** (22 mg, 40 µmol) in the mixture of THF (2 mL) and ethanol (0.5 mL), and the reaction mixture was stirred at rt for 5 h. Acetic acid (50 µL) was then added, the organic solvents were removed on a rotary evaporator, and the product was isolated by preparative HPLC (Interchim Uptisphere Strategy PhC4 250×21.2 mm 5 µm, solvent flow rate 18 mL/min, gradient 30% to 80% A:B, A – acetonitrile + 0.1% (v/v) $HCO_2H$, B – water + 0.1% (v/v) $HCO_2H$) and freeze-dried from dioxane to give 15 mg (72%) of **3** as orange-brown solid (mixture of (*E/Z*)-stereoisomers in ratio 0.65:0.35).

$^1$H NMR (400 MHz, DMSO-$d_6$): δ 12.00 (s, 1H, major isomer+minor isomer), 8.45 (s, 0.35H, minor isomer), 7.63 (s, 0.65H, major isomer), 7.54 (d, *J* = 8.6 Hz, 0.65H, major isomer), 7.52 (d, *J* = 8.4 Hz, 0.35H, minor isomer), 7.20 (dd, *J* = 17.5, 11.0 Hz, 0.35H, minor isomer), 7.01 (d, *J* = 2.7 Hz, 1H, major isomer + minor isomer), 6.97 (d, *J* = 2.6 Hz, 0.65H, major isomer), 6.89 – 6.85 (m, 1.35H, major isomer + 2×minor isomer), 6.82 (dd, *J* = 8.7, 2.8 Hz, 0.35H, minor isomer), 6.80 – 6.75 (m, 1.30H, 2×major isomer), 6.49 – 6.38 (m, 1H, major isomer + minor isomer), 5.91 (dd, *J* = 17.5, 1.1 Hz, 0.65H, major isomer), 5.66 (dd, *J* = 17.4, 1.6 Hz, 0.35H, minor isomer), 5.29 (dd, *J* = 10.9, 1.1 Hz, 0.65H, major isomer), 5.20 (dd, *J* = 10.9, 1.6 Hz, 0.35H, minor isomer), 3.16 – 3.02 (m, 2H, major isomer + minor isomer), 3.01 (s, 3.90H, major isomer), 2.98 (s, 2.10H, minor isomer), 2.97 (s, 2.10H, minor isomer), 2.93 (s, 3.90H, major isomer), 2.23 – 2.15 (m, 2H, major isomer + minor isomer), 1.56 – 1.38 (m, 4H, major isomer + minor isomer), 1.33 – 1.19 (m, 2H, major isomer + minor isomer), 0.61 (s, 3H, major isomer + minor isomer), 0.17 (s, 3H, major isomer + minor isomer).

$^{13}C$ NMR (101 MHz, DMSO-$d_6$): δ 174.5, 155.4, 154.9, 149.7, 149.4, 149.3, 149.0, 146.1, 144.4, 138.8, 138.5, 138.2, 137.1, 135.80, 135.76, 135.1, 134.0, 133.9, 131.4, 127.2, 126.6, 125.2, 124.4, 117.0, 116.3, 116.0, 115.7, 115.1, 113.2, 113.0, 112.3, 110.8, 108.6, 33.6, 30.4, 29.6, 29.5, 25.9, 24.3, -0.4, -5.4.
HRMS ($C_{28}H_{39}N_5O_3Si$): *m/z* (positive mode) = 522.2892 (found $[M+H]^+$), 522.2895 (calc.).

#### Synthesis of photoactivatable fluorescent probes

##### 1-NHS

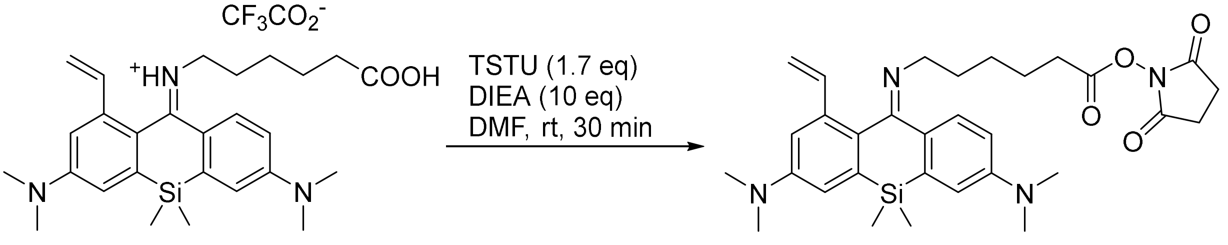

**1** **1-NHS**

In an amber vial, compound **1** (24 mg, 0.055 mmol) and *O*-(*N*-succinimidyl)-*N*,*N*,*N'*,*N'*-tetramethyluronium tetrafluoroborate (TSTU; 28 mg, 0.093 mmol, 1.7 equiv) were dissolved in DMF (500 µL). Ethyldiisopropylamine (DIEA; 96 µL, 0.55 mmol) was added, and the reaction mixture was stirred for 30 min at rt. The volatiles were removed *in vacuo*. The product was isolated by flash chromatography on a Biotage Isolera system (12g Interchim SiHP 30 µm cartridge, gradient 0% to 100% A/B, A = 10% MeOH – $CH_2Cl_2$, B = $CH_2Cl_2$). The product eluted at 50% A/B was freeze-dried from dioxane to yield 21 mg (68%) of **1-NHS** as an orange solid.

HRMS ($C_{31}H_{40}N_4O_4Si$): *m/z* (positive mode) = 561.2886 (found $[M+H]^+$), 561.2892 (calc.).

##### 2-NHS

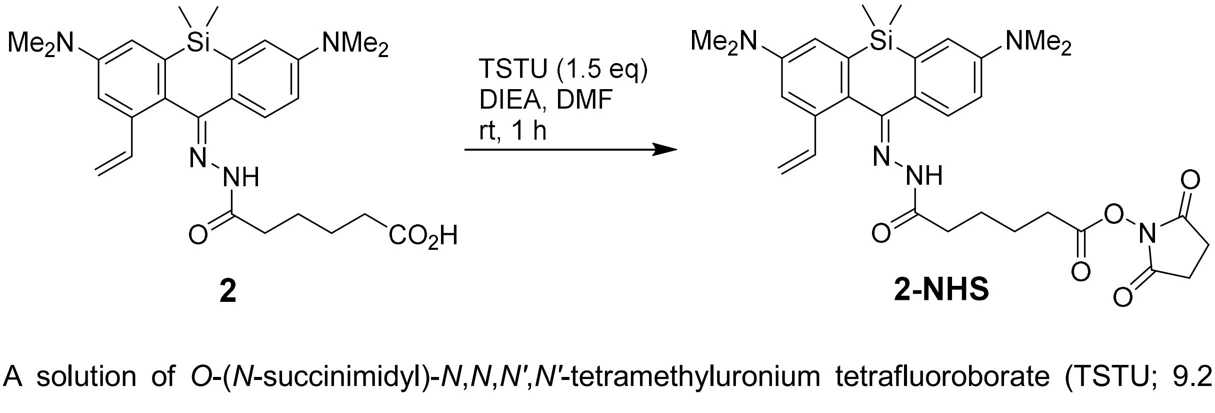

**2** **2-NHS**

A solution of *O*-(*N*-succinimidyl)-*N*,*N*,*N'*,*N'*-tetramethyluronium tetrafluoroborate (TSTU; 9.2 mg, 30.4 µmol, 1.5 equiv.) in DMF (100 µL) was added to the solution of **2** (10 mg, 20.3 µmol) in the mixture of DMF (100 µL) and ethyldiisopropylamine (DIEA; 50 µL), and the reaction mixture was stirred at rt for 1 h. The organic solvents were removed *in vacuo*, and the product was isolated by flash column chromatography (12 g Interchim SiHP 30 µm cartridge, gradient 0% to 50% EtOAc/$CH_2Cl_2$) and freeze-dried from 1,4-dioxane to yield 10.6 mg (89%) of **2-NHS** as light cream-colored solid as a mixture of (*E/Z*)-stereoisomers (ratio 0.64:0.36; each existing in the form of a pair of *(E/Z)* amide bond rotamers, of which the least abundant is very minor (<10%) and its signals in the $^1$H NMR data are ignored).

$^1$H NMR (400 MHz, $CD_3CN$): δ 9.24 (s, 0.08H, minor isomer/minor rotamer), 8.85 (s, 0.23H, major isomer/minor rotamer), 8.69 (s, 0.28H, minor isomer/major rotamer), 8.23 (s, 0.40H, major isomer/major rotamer), 7.58 (d, J = 8.5 Hz, 0.40H, major isomer/major rotamer), 7.55 (s, 0.28H, minor isomer/major rotamer), 7.46 (d, J = 8.6 Hz, 0.28H, minor isomer/major rotamer), 7.27 (dd, J = 17.4, 10.9 Hz, 0.23H, major isomer/minor rotamer), 7.08 – 7.03 (m, 1H), 6.99 – 6.91 (m, 2H), 6.86 – 6.77 (m, 1H), 6.58 – 6.45 (m, 0.63H, major isomer/major rotamer + minor isomer/major rotamer), 5.89 (dd, *J* = 17.4, 1.1 Hz, 0.23H, major isomer/minor rotamer), 5.87 (dd, *J* = 17.4, 1.1 Hz, 0.40H, major isomer/major rotamer), 5.68 (dd, *J* = 17.5, 1.5 Hz, 0.28H, minor isomer/major rotamer), 5.28 (dd, *J* = 10.9, 1.1 Hz, 0.23H, major isomer/minor rotamer), 5.27 (dd, *J* = 11.0, 1.1 Hz, 0.40H, major isomer/major rotamer), 5.22 (dd, *J* = 11.0, 1.5 Hz, 0.28H, minor isomer/major rotamer), 2.87 – 2.51 (m, 2H), 2.20 – 2.07 (m, 2H + $H_2O$), 1.82 – 1.60 (m, 4H), 0.63 (s, 1.08H, minor isomer/major+minor rotamer), 0.62 (s, 1.89H, major isomer/major+minor rotamer), 0.22 (s, 1.08H, minor isomer/major+minor rotamer), 0.21 (s, 1.89H, major isomer/major+minor rotamer).

$^{13}$C NMR (101 MHz, $CD_3CN$): δ 139.2, 136.8, 128.3, 127.6, 117.9, 116.6, 114.3, 113.8, 113.7, 113.3, 111.8, 110.1, 40.8, 40.5, 35.0, 32.9, 31.3, 26.4, 25.0, -0.4, -5.1 (indirect detection from a gHSQC experiment, only H-coupled carbons are resolved).

HRMS ($C_{31}H_{39}N_5O_5Si$): *m/z* (positive mode) = 590.2790 (found $[M+H]^+$), 590.2793 (calc.).

**3-NHS**

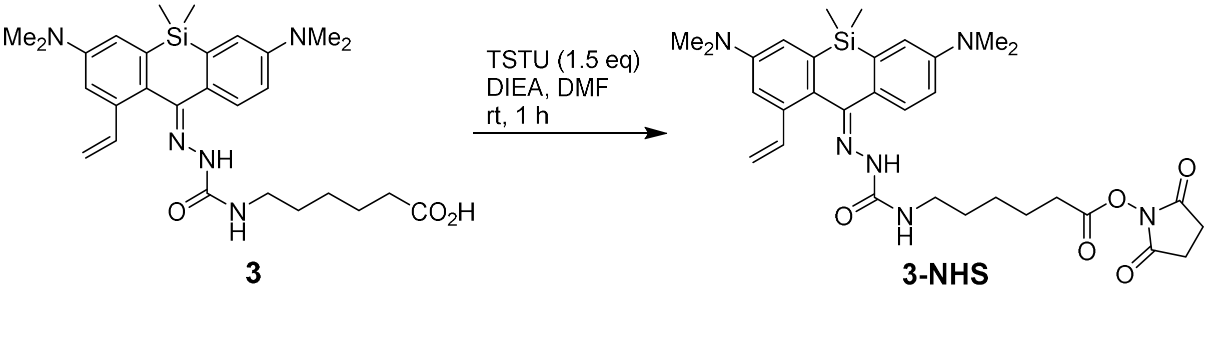

A solution of *O*-(*N*-succinimidyl)-*N*,*N*,*N'*,*N'*-tetramethyluronium tetrafluoroborate (TSTU; 8.6 mg, 28.7 µmol, 1.5 equiv.) in DMF (100 µL) was added to the solution of **3** (10 mg, 19.2 µmol)

in the mixture of DMF (100 µL) and ethyldiisopropylamine (DIEA; 50 µL), and the reaction mixture was stirred at rt for 1 h. The organic solvents were removed *in vacuo*, and the product was isolated by flash column chromatography (12 g Interchim SiHP 30 µm cartridge, gradient 0% to 80% EtOAc/$CH_2Cl_2$) and freeze-dried from 1,4-dioxane to yield 9.5 mg (80%) of **3-NHS** as white solid (mixture of (*E/Z*)-stereoisomers in ratio 0.58:0.42).

$^1$H NMR (400 MHz, $CDCl_3$): δ 8.21 (s, 0.42H, minor isomer), 7.78 (s, 0.42H, minor isomer), 7.76 (s, 0.58H, major isomer), 7.68 (s, 0.58H, major isomer), 7.62 – 7.52 (m, 1H, major isomer + minor isomer), 7.22 (dd, *J* = 17.4, 10.9 Hz, 0.42H, minor isomer), 7.14 – 6.80 (m, 3H, 3× major isomer + 3× minor isomer), 6.55 (dd, J = 17.4, 10.9 Hz, 0.58H, major isomer), 6.27 (t, *J* = 5.9 Hz, 0.42H, minor isomer), 6.20 (t, *J* = 6.0 Hz, 0.58H, major isomer), 5.84 (d, J = 17.4 Hz, 0.58H, major isomer), 5.71 (d, J = 17.4 Hz, 0.42H, minor isomer), 5.39 (d, *J* = 10.9 Hz, 0.58H, major isomer), 5.35 (d, *J* = 10.6 Hz, 0.42H, minor isomer), 3.42 – 3.23 (m, 2H, 2× major isomer + 2× minor isomer), 3.07 (s, 9.48H, 12× major isomer + 6× minor isomer), 3.05 (s, 2.52H, 6× minor isomer), 2.84 (s, 4H, 4× major isomer + 4× minor isomer), 2.65 – 2.58 (m, 2H, 2× major isomer + 2× minor isomer), 1.79 (p, *J* = 7.4 Hz, 2H, 2× major isomer + 2× minor isomer), 1.65 – 1.45 (m, 4H, 4× major isomer + 4× minor isomer), 0.69 (s, 3H, 3× major isomer + 3× minor isomer), 0.29 (s, 3H, 3× major isomer + 3× minor isomer).
HRMS ($C_{32}H_{42}N_6O_5Si$): *m/z* (positive mode) = 619.3049 (found $[M+H]^+$), 619.3059 (calc.).

**3-Halo**

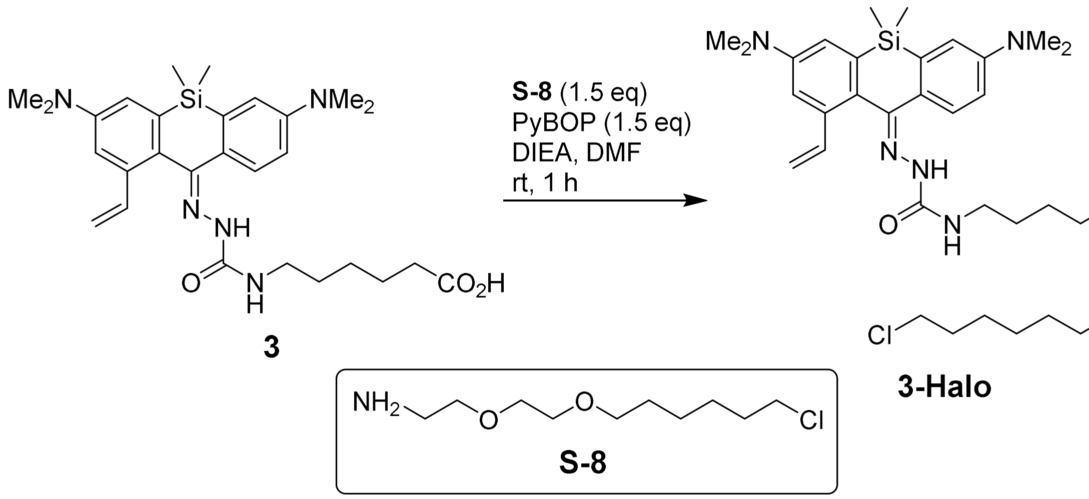

A solution of (benzotriazol-1-yloxy)tripyrrolidinophosphonium hexafluorophosphate (PyBOP; 7.5 mg, 14.4 µmol, 1.5 equiv.) in DMF (50 µL) was added to the solution of **3** (5 mg, 9.58 µmol) and **S-8** (HaloTag(O2) amine [13]; 3.2 mg, 14.4 µmol, 1.5 equiv.) in the mixture of DMF (150 µL) and ethyldiisopropylamine (DIEA; 40 µL), and the reaction mixture was stirred at rt for 1 h. The organic solvents were removed *in vacuo*, and the product was isolated by preparative

HPLC (Interchim Uptisphere Strategy PhC4 250×21.2 mm 5 µm, solvent flow rate 18 mL/min, gradient 40% to 80% A:B, A – acetonitrile + 0.1% (v/v) $HCO_2H$, B – water + 0.1% (v/v) $HCO_2H$). An attempt to freeze-dry the sample from dioxane gave 6 mg (86%) of **3-Halo** as viscous yellow oil which does not solidify (mixture of (*E/Z*)-stereoisomers in ratio 0.60:0.40).

$^1$H NMR (400 MHz, $CD_3CN$): δ 7.99 (s, 0.40H, minor isomer), 7.58 (d, *J* = 8.3 Hz, 0.60H, major isomer), 7.56 (d, *J* = 8.2 Hz, 0.40H, minor isomer), 7.39 (s, 0.60H, major isomer), 7.25 (dd, J = 17.5, 10.9 Hz, 0.40H, minor isomer), 7.06 (d, J = 2.8 Hz, 0.40H, minor isomer), 7.04 (d, J = 2.6 Hz, 0.60H, major isomer), 6.98 (d, J = 2.7 Hz, 0.60H, major isomer), 6.95 – 6.89 (m, 1.40H, 2× major isomer + minor isomer), 6.83 (dd, J = 8.7, 2.8 Hz, 0.40H, minor isomer), 6.80 (dd, J = 8.6, 2.8 Hz, 0.60H, major isomer), 6.54 (dd, J = 17.5, 11.0 Hz, 0.60H, major isomer), 6.39 (t, J = 5.7 Hz, 0.60H, major isomer), 6.29 (t, J = 5.9 Hz, 0.40H, minor isomer), 5.89 (dd, J = 17.5, 1.1 Hz, 0.60H, major isomer), 5.65 (dd, J = 17.5, 1.6 Hz, 0.40H, minor isomer), 5.28 (dd, J = 11.0, 1.0 Hz, 0.60H, major isomer), 5.22 (dd, J = 11.0, 1.6 Hz, 0.40H, minor isomer), 3.561 (t, J = 6.7 Hz, 1.20H, 2× major isomer), 3.556 (t, J = 6.7 Hz, 0.80H, 2× minor isomer), 3.52 – 3.46 (m, 4H, 4× major isomer + 4× minor isomer), 3.46 – 3.37 (m, 4H, 4× major isomer + 4× minor isomer), 3.29 – 3.23 (m, 2H, 2× major isomer + 2× minor isomer), 3.22 – 3.12 (m, 2H, 2× major isomer + 2× minor isomer), 3.03 (s, 3.60H, 6× major isomer), 3.00 (s, 2.40H, 6× minor isomer), 2.99 (s, 2.40H, 6× minor isomer), 2.96 (s, 3.60H, 6× major isomer), 2.11 (t, J = 7.5 Hz, 0.80H, 2× minor isomer), 2.11 (t, J = 7.5 Hz, 1.20H, 2× major isomer), 1.78 – 1.68 (m, 2H, 2× major isomer + 2× minor isomer), 1.63 – 1.45 (m, 6H, 6× major isomer + 6× minor isomer), 1.45 – 1.25 (m, 6H, 6× major isomer + 6× minor isomer), 0.62 (s, 1.80H, 3× major isomer), 0.42 (br.s, 2.40H, 6× minor isomer), 0.20 (s, 1.80H, 3× major isomer).
$^{13}$C NMR (101 MHz, $CD_3CN$): δ 140.1, 136.7, 128.0, 127.7, 117.8, 116.6, 116.5, 116.4, 114.3, 113.9, 113.6, 113.4, 112.5, 109.9, 71.7, 71.0, 70.4, 46.3, 40.8, 40.6, 40.2, 39.9, 36.9, 33.5, 30.7, 30.4, 27.4, 26.6, 26.4, -0.3, -5.0 (indirect detection from a gHSQC experiment, only H-coupled carbons are resolved).
HRMS ($C_{38}H_{59}ClN_6O_4Si$): *m/z* (positive mode) = 727.4124 (found $[M+H]^+$), 727.4128 (calc.).

## 3-BG

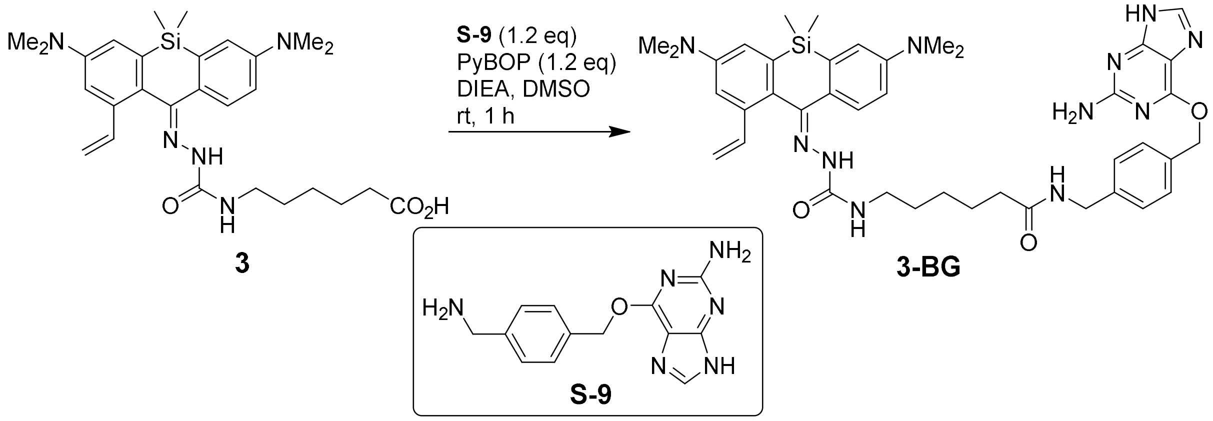

A solution of (benzotriazol-1-yloxy)tripyrrolidinophosphonium hexafluorophosphate (PyBOP; 6.2 mg, 12 µmol, 1.2 equiv.) in DMSO (30 µL) was added to the mixture of **3** (5.2 mg, 10 µmol), **S-9** (BG-$NH_2$ [14]; 3.2 mg, 12 µmol, 1.2 equiv.), ethyldiisopropylamine (DIEA; 40 µL) and DMSO (100 µL), and the reaction mixture was stirred at rt for 1 h. Excess DIEA was then removed *in vacuo*, and the product was isolated by preparative HPLC (Interchim Uptisphere Strategy PhC4 250×21.2 mm 5 µm, solvent flow rate 18 mL/min, gradient 30% to 70% A:B, A – acetonitrile + 0.1% (v/v) $HCO_2H$, B – water + 0.1% (v/v) $HCO_2H$) and freeze-dried from 1,4-dioxane to yield 7 mg (90%) of **3-BG** as white solid (mixture of (*E/Z*)-stereoisomers in ratio 0.70:0.30).

$^1$H NMR (400 MHz, $CD_3CN$ + 1%(v/v) TFA-*d*): δ 8.70 (s, 0.30H, minor isomer), 8.25 (s, 0.70H, major isomer), 8.08 (s, 0.70H, major isomer), 8.04 (s, 0.30H, minor isomer), 7.85 (d, J = 8.4 Hz, 0.70H, major isomer), 7.77 – 7.71 (m, 1.30H, major isomer + 2× minor isomer), 7.66 (dd, J = 4.5, 2.5 Hz, 0.30H, minor isomer), 7.63 – 7.56 (m, 2.30H, 2× major isomer + 3× minor isomer), 7.48 – 7.40 (m, 1.70H, 2× major isomer + minor isomer), 7.37 (dd, J = 8.5, 2.6 Hz, 0.30H, minor isomer), 7.32 – 7.26 (m, 2H, 2× major isomer + 2× minor isomer), 7.26 – 7.17 (m, 1H, major isomer + minor isomer), 6.52 (dd, J = 17.4, 11.0 Hz, 0.70H, major isomer), 5.86 (d, J = 17.4 Hz, 0.70H, major isomer), 5.76 (d, J = 17.4 Hz, 0.30H, minor isomer), 5.55 – 5.49 (m, 1.30H, major isomer + 2× minor isomer), 5.44 – 5.36 (m, 1.70H, 2× major isomer + minor isomer), 4.41 – 4.26 (m, 2H, 2× major isomer + 2× minor isomer), 3.23 (s, 1.80H, 6× minor isomer), 3.25 – 3.11 (m, 2H, 2× major isomer + 2× minor isomer), 3.21 (s, 4.20H, 6× major isomer), 3.18 (s, 4.20H, 6× major isomer), 3.15 (s, 1.80H, 6× minor isomer), 2.28 – 2.20 (m, 2H, 2× major isomer + 2× minor isomer), 1.68 – 1.55 (m, 2H, 2× major isomer + 2× minor isomer), 1.55 – 1.44 (m, 2H, 2× major isomer + 2× minor isomer), 1.35 – 1.24 (m, 4H, 4× major isomer + 4× minor isomer), 0.72 (s, 3H, 3× major isomer + 3× minor isomer), 0.25 (s, 3H, 3× major isomer + 3× minor isomer).

$^{13}C$ NMR (101 MHz, $CD_3CN$ + 1%(v/v) TFA-*d*): δ 143.6, 136.8, 134.9, 129.9, 129.6, 128.9, 128.7, 125.1, 124.4, 123.7, 122.8, 120.8, 120.2, 119.3, 117.4, 117.3, 70.7, 47.1, 45.2, 43.5, 40.5, 36.5, 30.4, 26.9, 26.3, -1.0, -5.5.
HRMS ($C_{41}H_{51}N_{11}O_3Si$): *m/z* (positive mode) = 387.7045 (found $[M+2H]^{2+}$), 387.7046 (calc.).

#### Compound S-10

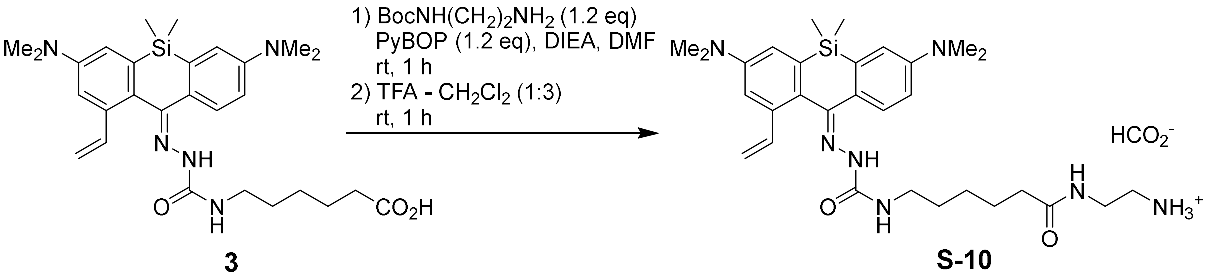

A solution of (benzotriazol-1-yloxy)tripyrrolidinophosphonium hexafluorophosphate (PyBOP; 8.6 mg, 16.6 µmol, 1.2 equiv.) in DMF (40 µL) was added to the solution of **3** (7.2 mg, 13.8 µmol), *N*-Boc-ethylenediamine (2.7 mg, 16.6 µmol, 1.2 equiv.) in the mixture of DMF (100 µL) and ethyldiisopropylamine (DIEA; 50 µL), and the reaction mixture was stirred at rt for 1 h. The organic solvents were removed *in vacuo*, the residue was redissolved in the mixture of trifluoroacetic acid (50 µL) and $CH_2Cl_2$ (150 µL) and stirred at rt for 1 h. The solvents were then evaporated, the product was isolated by preparative HPLC (Interchim Uptisphere Strategy PhC4 250×21.2 mm 5 µm, solvent flow rate 18 mL/min, gradient 20% to 70% A:B, A – acetonitrile + 0.1% (v/v) $HCO_2H$, B – water + 0.1% (v/v) $HCO_2H$) and freeze-dried from 1,4-dioxane to give 7.3 mg (87%) of **S-10** as light pink solid (mixture of (*E/Z*)-stereoisomers in ratio 0.70:0.30).

$^1H$ NMR (400 MHz, $CD_3CN$): δ 8.35 (br.s, 0.70H, major isomer), 8.20 (br.s, 0.30H, minor isomer), 7.62 (br.s, 0.30H, minor isomer), 7.60 (br.s, 0.70H, major isomer), 7.57 (d, *J* = 8.5 Hz, 0.70H, major isomer), 7.54 (d, *J* = 8.6 Hz, 0.30H, minor isomer), 7.23 (dd, J = 17.4, 10.9 Hz, 0.30H, minor isomer), 7.03 (d, *J* = 2.7 Hz, 0.30H, minor isomer), 7.02 (d, *J* = 2.6 Hz, 0.70H, major isomer), 6.95 (d, J = 2.6 Hz, 0.70H, major isomer), 6.93 – 6.88 (m, 1.30H, major isomer + 2× minor isomer), 6.81 – 6.74 (m, 1H, major isomer + minor isomer), 6.53 (dd, J = 17.4, 11.0 Hz, 0.70H, major isomer), 6.46 (t, J = 6.0 Hz, 0.70H, major isomer), 6.34 (t, J = 6.0 Hz, 0.30H, minor isomer), 5.86 (d, *J* = 17.5 Hz, 0.70H, major isomer), 5.64 (dd, *J* = 17.4, 1.6 Hz, 0.30H, minor isomer), 5.25 (d, *J* = 11.1 Hz, 0.70H, major isomer), 5.19 (dd, *J* = 10.9, 1.6 Hz, 0.30H, minor isomer), 3.29 (br.s, 2H, 2× major isomer + 2× minor isomer), 3.20 – 3.09 (m, 2H, 2× major isomer + 2× minor isomer), 3.00 (s, 4.20H, 6× major isomer), 2.97 (s, 3.60H, 12× minor isomer), 2.94 (s, 4.20H, 6× major isomer), 2.86 (br.s, 2H, 2× major isomer + 2× minor isomer), 2.15 – 2.05 (m, 2H, 2× major isomer + 2× minor isomer), 1.60 – 1.41 (m, 4H, 4× major isomer + 4× minor isomer), 1.35 – 1.20 (m, 2H, 2× major isomer + 2× minor isomer), 0.61 (s, 0.90H, 3× minor isomer), 0.60 (s, 2.10H, 3× major isomer), 0.20 (s, 0.90H, 3× minor isomer), 0.19 (s, 2.10H, 3× major isomer).

$^{13}$C NMR (101 MHz, $CD_3CN$): δ 139.5, 136.1, 127.7, 127.1, 117.3, 115.9, 115.8, 113.6, 113.0, 111.8, 109.4, 40.3, 40.1, 39.9, 39.5, 38.5, 36.0, 30.2, 26.4, 25.4, -0.9, -5.7 (indirect detection from a gHSQC experiment, only H-coupled carbons are resolved).

HRMS ($C_{30}H_{45}N_7O_2Si$): *m/z* (positive mode) = 564.3477 (found $[M+H]^+$), 564.3477 (calc.).

#### 3-Lyso

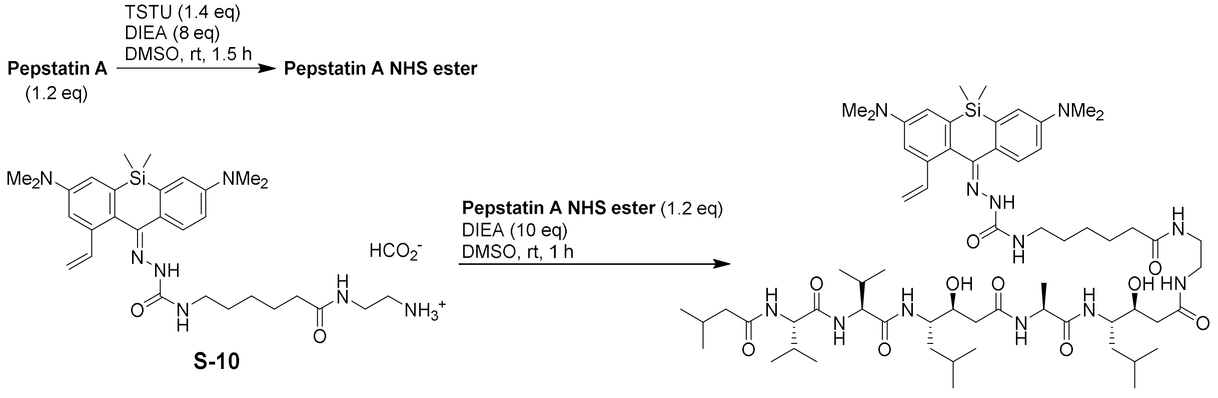

**3-Lyso**

Pepstatin A NHS ester solution was prepared as described in [15]; specifically, a solution of *O*-(*N*-succinimidyl)-*N*,*N*,*N'*,*N'*-tetramethyluronium tetrafluoroborate (TSTU; 5 mg, 16.7 µmol, 1.2 equiv. relative to Pepstatin A or 1.4 equiv. relative to **S-10**) in DMSO (50 µL) was added to a mixture of Pepstatin A (9.9 mg, 14.4 µmol, 1.2 equiv.) and DIEA (25 µL, 0.12 mmol, ~8 equiv.) in anhydrous DMSO (500 µL) and the mixture was stirred at rt for 1.5 h. TLC control (silica, 10% methanol – $CH_2Cl_2$) showed complete conversion: $R_f$ = 0.06 (starting material), 0.32 (product), both stained light-brown with vanillin stain.

The resulting solution of Pepstatin A NHS ester was added to the solution of **S-10** (formate salt; 7.3 mg, ~12 µmol) in the mixture of DIEA (35 µL) and DMSO (150 µL), and the resulting mixture was stirred at rt for 1 h. The organic solvents were removed *in vacuo* overnight, the residue was dissolved in minimal volume of DMSO, the product was isolated by preparative HPLC (Interchim Uptisphere Strategy PhC4 250×21.2 mm 5 µm, solvent flow rate 18 mL/min, gradient 30% to 80% A:B, A – acetonitrile + 0.1% (v/v) $HCO_2H$, B – water + 0.1% (v/v) $HCO_2H$) and freeze-dried from aqueous 1,4-dioxane to give 7.6 mg (51%) of **3-Lyso** as white solid (mixture of (*E/Z*)-stereoisomers in ratio 0.67:0.33).

$^1H$ NMR (400 MHz, DMSO-$d_6$): δ 8.45 (s, 0.33H, minor isomer), 7.94 (d, J = 7.2 Hz, 1H, major isomer + minor isomer), 7.85 – 7.70 (m, 3.33H, 3× major isomer + 4× minor isomer), 7.63 (s, 0.67H, major isomer), 7.54 (d, J = 8.6 Hz, 0.67H, major isomer), 7.51 (d, *J* = 8.7 Hz, 0.33H, minor isomer), 7.48 (d, *J* = 8.8 Hz, 1H, major isomer + minor isomer), 7.34 (d, J = 9.2 Hz, 1H, major isomer + minor isomer), 7.20 (dd, J = 17.4, 10.9 Hz, 0.33H, minor isomer), 7.01 (d, J = 2.7 Hz, 1H, major isomer + minor isomer), 6.96 (d, J = 2.6 Hz, 0.67H, major isomer), 6.90 – 6.84 (m, 1.33H, major isomer + 2× minor isomer), 6.82 (dd, *J* = 8.7, 2.8 Hz, 0.33H, minor isomer), 6.80 – 6.74 (m, 1.34H, 2× major isomer), 6.48 – 6.38 (m, 1H, major isomer + minor isomer), 5.91 (dd, *J* = 17.5, 1.1 Hz, 0.67H, major isomer), 5.65 (dd, *J* = 17.4, 1.6 Hz, 0.33H, minor isomer), 5.29 (dd, *J* = 11.0, 1.1 Hz, 0.67H, major isomer), 5.19 (dd, *J* = 11.0, 1.6 Hz, 0.33H, minor isomer), 4.89 – 4.80 (m, 2H, 2× major isomer + 2× minor isomer), 4.25 (p, *J* = 7.0 Hz, 1H, major isomer + minor isomer), 4.19 (dd, *J* = 8.6, 7.5 Hz, 1H, major isomer + minor isomer), 4.13 (dd, *J* = 9.0, 7.2 Hz, 1H, major isomer + minor isomer), 3.88 – 3.73 (m, 4H, 4× major isomer + 4× minor isomer), 3.20 – 3.03 (m, 5H, 5× major isomer + 5× minor isomer), 3.01 (s, 4.02H, 6× major isomer), 2.98 (s, 1.98H, 6× minor isomer), 2.97 (s, 1.98H, 6× minor isomer), 2.93 (s, 4.02H, 6× major isomer), 2.17 – 2.00 (m, 6H, 6× major isomer + 6× minor isomer), 2.00 – 1.88 (m, 3H, 3× major isomer + 3× minor isomer), 1.59 – 1.30 (m, 8H, 8× major isomer + 8× minor isomer), 1.29 – 1.16 (m, 8H, 8× major isomer + 8× minor isomer), 0.91 – 0.75 (m, 30H, 30× major isomer + 30× minor isomer), 0.61 (s, 3H, 3× major isomer + 3× minor isomer), 0.17 (s, 3H, 3× major isomer + 3× minor isomer).
$^{13}C$ NMR (101 MHz, DMSO-$d_6$): δ 138.2, 135.0, 126.6, 127.1, 124.9, 116.7, 116.0, 115.1, 113.2, 113.0, 112.3, 110.7, 108.5, 68.9, 57.9, 57.7, 50.6, 50.1, 48.4, 44.3, 40.0, 39.9, 39.8, 39.6, 39.1, 38.5, 35.3, 30.2, 29.6, 28.7, 26.0, 25.6, 24.9, 24.1, 22.7, 21.6, 18.6, 18.1, -0.6, -5.5.
HRMS ($C_{64}H_{106}N_{12}O_{10}Si$): *m/z* (positive mode) = 1231.7991 (found $[M+H]^+$), 1231.7997 (calc.).

## 3-Tz

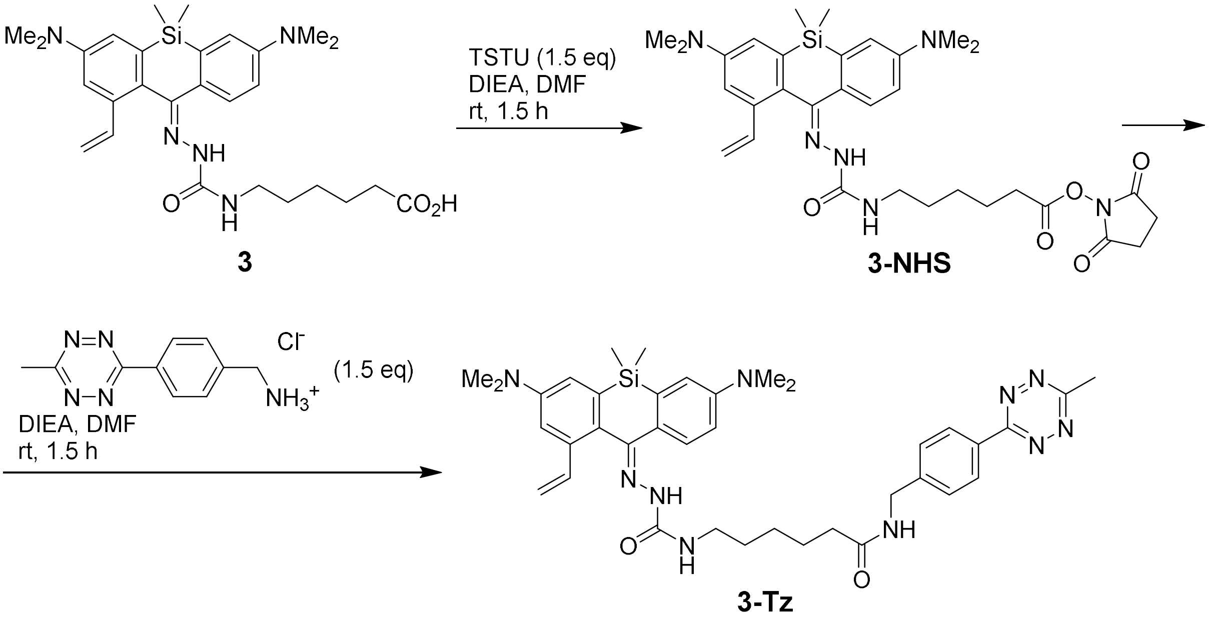

A solution of *O*-(*N*-succinimidyl)-*N*,*N*,*N'*,*N'*-tetramethyluronium tetrafluoroborate (TSTU; 6.9 mg, 23 µmol, 1.5 equiv.) in DMF (50 µL) was added to the solution of **3** (8 mg, 15.3 µmol) in the mixture of DMF (100 µL) and ethyldiisopropylamine (DIEA; 30 µL), and the reaction mixture was stirred at rt for 1.5 h. The resulting solution of **3-NHS** was added to the stirred suspension of 4-(6-methyl-1,2,4,5-tetrazin-3-yl)benzylamine hydrochloride (4.6 mg, 23 µmol, 1.5 equiv.) in the mixture of DMF (70 µL) and DIEA (50 µL), and the reaction mixture was stirred at rt for 1.5 h. The organic solvents were removed *in vacuo*, the residue was dissolved in minimal volume of DMSO, the product was isolated by preparative HPLC (Interchim Uptisphere Strategy PhC4 250×21.2 mm 5 µm, solvent flow rate 18 mL/min, gradient 40% to 80% A:B, A – acetonitrile + 0.1% (v/v) $HCO_2H$, B – water + 0.1% (v/v) $HCO_2H$) and freeze-dried from 1,4-dioxane to give 8.6 mg (80%) of **3-Tz** as pink solid (mixture of (*E/Z*)-stereoisomers in ratio 0.60:0.40).

$^1$H NMR (400 MHz, $CDCl_3$): δ 8.57 – 8.51 (m, 2H, 2× major isomer + 2× minor isomer), 8.12 (br.s, 0.40H, minor isomer), 7.61 (br.s, 1.20H, 2× major isomer), 7.53 (d, J = 8.6 Hz, 0.40H, minor isomer), 7.51 – 7.45 (m, 2H, 2× major isomer + 2× minor isomer), 7.23 (dd, J = 17.4, 10.9 Hz, 0.40H, minor isomer), 7.02 – 6.84 (m, 2.60H, 3× major isomer + 2× minor isomer), 6.78 – 6.69 (m, 0.40H, minor isomer), 6.56 (dd, J = 17.4, 10.9 Hz, 0.60H, major isomer), 6.36 (t, J = 5.9 Hz, 0.40H, minor isomer), 6.26 (t, J = 6.0 Hz, 0.60H, major isomer), 6.15 – 6.04 (m, 1H), 5.79 (d, *J* = 17.4 Hz, 0.60H, major isomer), 5.63 (d, J = 17.4 Hz, 0.40H, minor isomer), 5.31 (d, *J* = 10.9 Hz, 0.60H, major isomer), 5.23 (d, J = 11.0 Hz, 0.40H, minor isomer), 4.54 (t, J = 5.7 Hz, 2H, 2× major isomer + 2× minor isomer), 3.29 (q, J = 6.8 Hz, 2H, 2× major isomer + 2× minor isomer), 3.09 (s, 2.40H, 6× minor isomer), 3.03 (s, 7.20H, 12× major isomer), 3.02 (s, 2.40H, 6× minor isomer), 2.32 – 2.24 (m, 2H, 2× major isomer + 2× minor isomer), 1.80 –

1.66 (m, 2H, 2× major isomer + 2× minor isomer), 1.64 – 1.51 (m, 2H, 2× major isomer + 2× minor isomer), 1.47 – 1.34 (m, 2H, 2× major isomer + 2× minor isomer).

$^{13}C$ NMR (101 MHz, $CDCl_3$): δ 138.8, 135.2, 128.3, 128.1, 127.3, 126.5, 116.8, 116.6, 116.4, 113.0, 112.4, 109.4, 43.0, 40.1, 39.2, 36.5, 29.7, 26.2, 25.1, 21.0, -0.5, -5.3 (indirect detection from a gHSQC experiment, only H-coupled carbons are resolved).

HRMS ($C_{38}H_{48}N_{10}O_2Si$): *m/z* (positive mode) = 705.3808 (found $[M+H]^+$), 705.3804 (calc.).

#### Supplementary references

[1] Walter, J. Spectral Unmixing for ImageJ – documentation (v1.2), https://imagej.nih.gov/ij/plugins/docs/SpectralUnmixing.pdf (accessed November 2022).

[2] Lincoln, R.; Bossi, M. L.; Remmel, M.; D'Este, E.; Butkevich, A. N.; Hell, S. W. A general design of caging-group-free photoactivatable fluorophores for live-cell nanoscopy. *Nat. Chem.* **2022**, *14*, 1013–1020.

[3] Uno, K.; Bossi, M. L.; Konen, T.; Belov, V. N.; Irie, M.; Hell, S. W. Asymmetric diarylethenes with oxidized 2-alkylbenzothiophen-3-yl units: chemistry, fluorescence, and photoswitching. *Adv. Optical Mater.* **2019**, *7*, 1801746.

[4] Ovesný, M.; Křížek, P.; Borkovec, J.; Švindrych, Z.; Hagen, G. M. ThunderSTORM: a comprehensive ImageJ plug-in for PALM and STORM data analysis and super-resolution imaging. *Bioinformatics* **2014**, *30*, 2389–2390.

[5] Butkevich, A. N.; Ta, H.; Ratz, M.; Stoldt, S.; Jakobs, S.; Belov, V. N.; Hell, S. W. Two-Color 810 nm STED Nanoscopy of Living Cells with Endogenous SNAP-Tagged Fusion Proteins. *ACS Chem. Biol.* **2018**, *13*, 475–480.

[6] Thevathasan, J. V.; Kahnwald, M.; Cieslinski, K.; Hoess, P.; Peneti, S. K.; Reitberger, M.; Heid, D.; Kasuba, K. C.; Hoerner, S. J.; Li, Y. M.; Wu, Y. L.; Mund, M.; Matti, U.; Pereira, P. M.; Henriques, R.; Nijmeijer, B.; Kueblbeck, M.; Jimenez Sabinina, V.; Ellenberg, J.; Ries, J. Nuclear pores as versatile reference standards for quantitative superresolution microscopy. *Nat. Methods* **2019**, *16*, 1045–1053.

[7] Wurm, C. A.; Neumann, D.; Lauterbach, M. A.; Harke, B.; Egner, A.; Hell, S. W.; Jakobs, S. Nanoscale distribution of mitochondrial import receptor Tom20 is adjusted to cellular conditions and exhibits an inner-cellular gradient. *Proc. Natl. Acad. Sci. U.S.A.* **2011**, *108*, 13546–13551.

[8] Xu, K.; Babcock, H. P.; Zhuang, X. Dual-objective STORM reveals three-dimensional filament organization in the actin cytoskeleton. *Nat. Methods* **2012**, *9*, 185–188.

[9] Lukinavičius, G.; Reymond, L.; D'Este, E.; Masharina, A.; Göttfert, F.; Ta, H.; Güther, A.; Fournier, M.; Rizzo, S.; Waldmann, H.; Blaukopf, C.; Sommer, C.; Gerlich, D. W.; Arndt, H.-D.; Hell, S. W.; Johnsson, K. Fluorogenic probes for live-cell imaging of the cytoskeleton. *Nat. Methods* **2014**, *11*, 731–733.

[10] Murrey, H. E.; Judkins, J. C.; am Ende, C. W.; Ballard, T. E.; Fang, Y.; Riccardi, K.; Di, L.; Guilmette, E. R.; Schwartz, J. W.; Fox, J. M.; Johnson, D. S. Systematic evaluation of

bioorthogonal reactions in live cells with clickable HaloTag ligands: implications for intracellular imaging. *J. Am. Chem. Soc.* **2015**, *137*, 11461–11475.

[11] Werther, P.; Yserentant, K.; Braun, K.; Kaltwasser, N.; Popp, C.; Baalmann, M.; Herten, D.-P.; Wombacher, R. Live-cell localization microscopy with a fluorogenic and self-blinking tetrazine probe. *Angew. Chem. Int. Ed.* **2020**, *59*, 804–810.

[12] Zou, L.; Braegelman, A. S.; Webber, M. J. Spatially defined drug targeting by in situ host-guest chemistry in a living animal. *ACS Cent. Sci.* **2019**, *5*, 1035–1043.

[13] Singh, V.; Wang, S.; Kool, E.T. Genetically encoded multispectral labeling of proteins with polyfluorophores on a DNA backbone. *J. Am. Chem. Soc.* **2013**, *135*, 6184–6191.

[14] Kobayashi, T.; Komatsu, T.; Kamiya, M.; Campos, C.; González-Gaitán, M.; Terai, T.; Hanaoka, K.; Nagano, T.; Urano, Y. Highly activatable and environment-insensitive optical highlighters for selective spatiotemporal imaging of target proteins. *J. Am. Chem. Soc.* **2012**, *134*, 11153–11160.

[15] Butkevich, A. N.; Lukinavičius, G.; D'Este, E.; Hell, S. W. Cell-permeant large Stokes shift dyes for transfection-free multicolor nanoscopy. *J. Am. Chem. Soc.* **2017**, *139*, 12378–12381.
